# Complex basis of hybrid female sterility and Haldane’s rule in *Heliconius* butterflies: Z-linkage and epistasis

**DOI:** 10.1101/2021.06.28.450252

**Authors:** Neil Rosser, Nathaniel B. Edelman, Lucie M. Queste, Michaela Nelson, Fernando Seixas, Kanchon K. Dasmahapatra, James Mallet

## Abstract

Hybrids between diverging populations are often sterile or inviable. Hybrid unfitness usually evolves first in the heterogametic sex – a pattern known as Haldane’s rule. The genetics of Haldane’s Rule have been extensively studied in species where the male is the heterogametic (XX/XY) sex, but its basis in taxa where the female is heterogametic (ZW/ZZ), such as Lepidoptera and birds, is largely unknown. Here, we analyse a new case of female hybrid sterility between geographic subspecies of *Heliconius pardalinus*. The two subspecies mate freely in captivity, but female F1 hybrids in both directions of cross are sterile. Sterility is due to arrested development of oocytes after they become differentiated from nurse cells, but before yolk deposition. We backcrossed fertile male F1 hybrids to parental females, and mapped quantitative trait loci (QTLs) for female sterility. We also identified genes differentially expressed in the ovary, and as a function of oocyte development. The Z chromosome has a major effect, similar to the “large X effect” in *Drosophila*, with strong epistatic interactions between loci at either end of the Z chromosome, and between the Z chromosome and autosomal loci on chromosomes 8 and 20. Among loci differentially expressed between females with arrested vs. non-arrested ovary development, we identified six candidate genes known also from *Drosophila melanogaster* and *Parage aegeria* oogenesis. This study is the first to characterize hybrid sterility using genome mapping in the Lepidoptera. We demonstrate that sterility is produced by multiple complex epistastic interactions often involving the sex chromosome, as predicted by the dominance theory of Haldane’s Rule.

## 1 Introduction

Hybrids between diverging populations are often sterile or inviable (Darwin, 1859; Presgraves, 2010). Because such examples of postzygotic incompatibility are common between species, elucidating their genetic basis is seen as key to understanding speciation (Nosil & Schluter, 2011; Butlin *et al*., 2012; Castillo & Barbash, 2017; Coughlan & Matute, 2020). Hybrid dysfunction often results from epistatic interactions among genes known as “Dobzhansky-Muller Incompatibilities” (Bateson, 1909; Dobzhansky, 1936; Muller, 1942; Coyne & Orr, 2004). Under the Dobzhansky-Muller model, diverging populations acquire different alleles at two or more loci. In hybrids, previously untested combinations of alleles at different loci are brought together and interact to reduce fitness (Orr, 1995; Brideau *et al*., 2006; Tang & Presgraves, 2009; Presgraves, 2007; Maheshwari & Barbash, 2011).

Dobzhansky-Muller incompatibilities (DMIs) may involve only a pair of genes (Sweigart *et al*., 2006), but they are perhaps more likely to be complex, even early in speciation (e.g. Phadnis, 2011; Kalirad & Azevedo, 2017). This is because the expected number of two-locus DMIs is predicted to increase approximately as the square of the number of divergent substitutions between species; the “snowball effect” (Orr, 1995; Orr & Turelli, 2001; Matute *et al*., 2010). Furthermore, DMIs involving more than two loci should accumulate even more rapidly, because, as the number of interacting loci increases, so too does the number of potentially negative combinations (Orr, 1995). In keeping with these predictions, widespread DMIs across the genomes of a number of species have been inferred from genetic association data (Good *et al*., 2008; Schumer *et al*., 2014). There is also evidence that polymorphic alleles with negative epistatic interactions are common even within species (Corbett-Detig *et al*., 2013)

One of the few generalisations about speciation is Haldane’s Rule, which states that among hybrids, when one sex is absent, rare, or sterile, it is usually the heterogametic sex (males in XX/XY systems and females in ZZ/ZW systems (Haldane, 1922). Greater sterility of the heterogametic sex has been found in 213 out of 223 pairs (*>*95%) of a diverse array of taxa, and has at least 10 phylogenetically independent origins (Schilthuizen *et al*., 2011; Delph & Demuth, 2016). The ubiquity of Haldane’s rule therefore suggests that postzygotic incompatibilities evolve with some predictability across a wide range of taxa (Coyne, 1992). Hybrid sterility of the heterogametic sex also evolves early, typically before hybrid inviability (Coyne & Orr, 1989a; 1997; Presgraves, 2010; 2002). It may therefore have a disproportionate role in reducing gene flow, and as such is of particular interest for understanding speciation (Ramsey *et al*., 2003; Coughlan & Matute, 2020).

Most explanations for Haldane’s rule depend on DMIs. The hypothesis to have received the most support is dominance theory, in which hybrid sterility and inviability are produced by interactions between the sex chromosomes and autosomes (Coyne & Orr, 2004). In the homogametic sex of hybrids, sex-linked alleles produce incompatibilities only if dominant, whereas in heterogametic hybrids both dominant and recessive sex-linked alleles can cause incompatibilities. If alleles causing incompatibilities are on average recessive, the heterogametic sex is expected to suffer more than the homogametic sex (Turelli & Orr, 1995; Orr, 1997; Turelli & Moyle, 2007). Nonetheless, male heterogametic species without strongly differentiated sex chromosomes also conform to Haldane’s Rule (Presgraves & Orr, 1998), suggesting that other forces also contribute, such as “faster-male” evolution (Wu & Davis, 1993) and faster evolution of the sex chromosome (Sackton *et al*., 2014). The genetic and molecular mechanisms of hybrid sterility have been identified in some cases (Brideau *et al*., 2006; Tang & Presgraves, 2009; Schartl, 2008; Mihola *et al*., 2009; Bayes & Malik, 2009), but this work has been primarily carried out in organisms with XX/XY sex determination, in which male hybrids are sterile or inviable.

Lepidoptera (butterflies and moths) yielded the first example of a sex linked trait (Doncaster & Raynor, 1906), even before *Drosophila* (Morgan, 1910; 1911). Lepidoptera are also among the groups of taxa Haldane considered when formulating his eponymous rule (Haldane, 1922). They have ZW/ZZ sex determination, where females are the heterogametic sex and, in accordance with Haldane’s Rule, are more susceptible to hybrid dysfunction (Presgraves, 2002). As such, they are critical in evaluating the relative impact of dominance and faster male evolution in Haldane’s rule, and have provided evidence that faster-Z evolution may contribute to the phenomenon in female heterogametic systems (Prowell Pashley, 1998; Sackton *et al*., 2014).

Several examples of Haldane’s Rule have been reported in *Heliconius* butterflies (Nymphalidae), which comprise about 48 species that occur throughout much of tropical America (Jiggins, 2017). Female hybrid sterility has been observed in crosses between *Heliconius cydno* (*sensu lato*) and *Heliconius melpomene* (Naisbit *et al*., 2002; Salazar *et al*., 2005; Śanchez *et al*., 2015), and also between geographically distant subspecies of *Heliconius melpomene* (Jiggins *et al*., 2001). Here, we investigate the genetic and molecular basis of Haldane’s rule in hybrids between two subspecies of *Heliconius pardalinus*: *H. pardalinus butleri* and *H. pardalinus sergestus*. These largely allopatric subspecies are strongly genetically differentiated, with *H. p. butleri* more closely related over most of its genome to its sympatric relative *Heliconius elevatus*, thereby rendering *H. pardalinus* paraphyletic (Rosser *et al*., 2019). They inhabit different habitats, with *H. p. sergestus* restricted to dry forests in the Huallaga/Mayo valleys of the Andes, and *H. p. butleri* inhabiting lowland rainforest across the adjacent Amazon basin (Fig. 1). Although they mate freely in captivity, they rarely co-occur in nature, and F1 hybrid females in both directions of cross are completely sterile (Rosser *et al*., 2019). Here, we characterize the ovary phenotype in parental populations, F1 hybrids and backcrosses. We use backcrosses to *H. p. butleri* to generate a QTL map and intersect these data with genes differentially expressed between fertile and sterile individuals, to identify candidate genes and epistatic interactions responsible for hybrid sterility.

**Figure 1:**
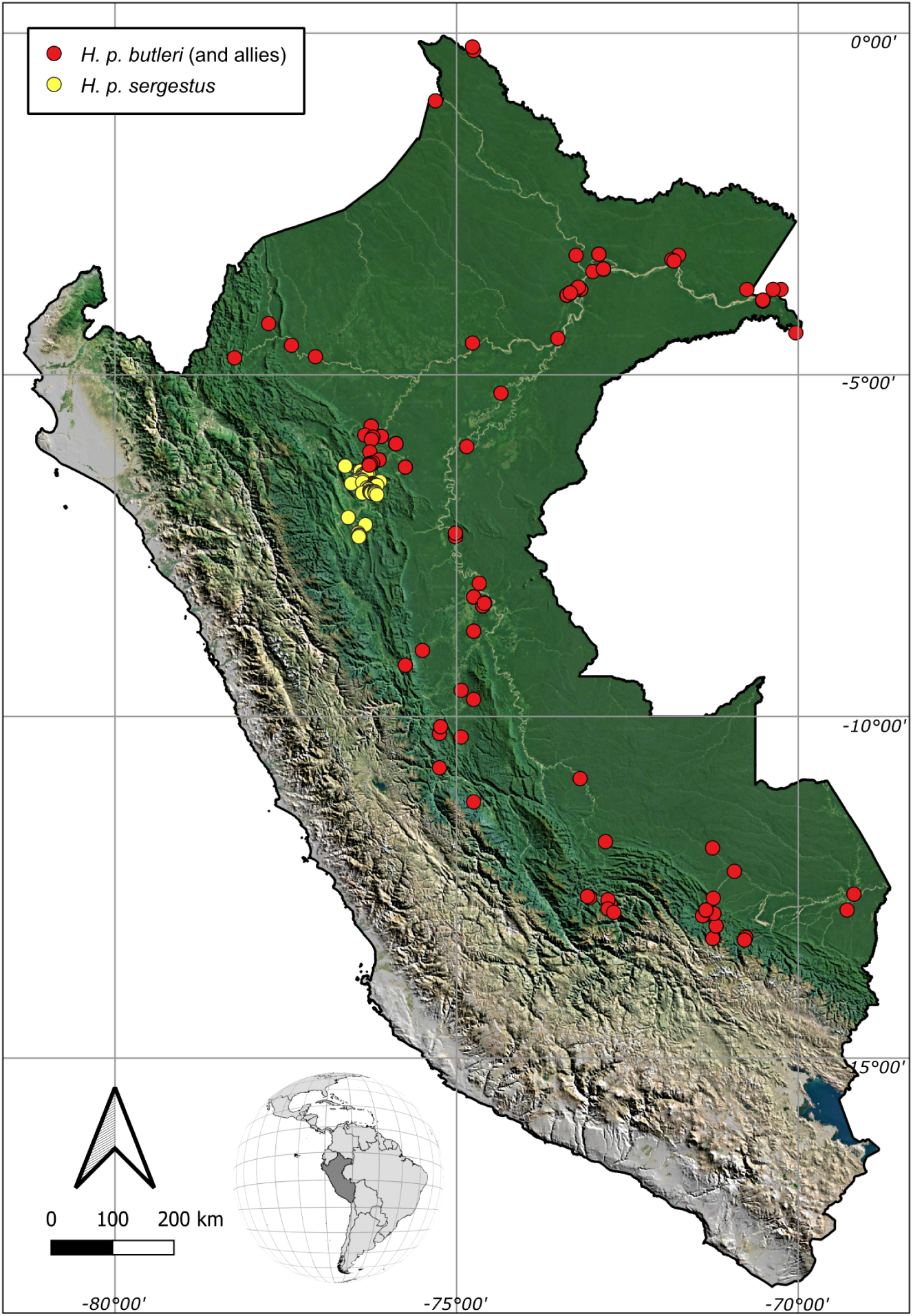
Distribution of *H. pardalinus* in Peru. The yellow dots correspond to collection localities of *H. pardalinus sergestus* and the red dots to *H. pardalinus butleri*, which intergrades with other subspecies of in central and southern Peru and the Amazon basin. Geographic data are from Rosser *et al*. (2019; 2012).

## 2 Materials and methods

### 2.1 Butterfly rearing, nucleic acid preservation and ovary dissection

Butterfly stocks were collected in the Departments of San Martín, Loreto and Ucayali, Peru, and captive populations of *H. p. sergestus* and *H. p. butleri* were established in insectaries in Tarapoto, Peru, as previously described (Rosser *et al*., 2019). Female butterflies were collected from insectaries 15 days after eclosion, allowing time for eggs to develop fully (Dunlap-Pianka *et al*., 1977; Naisbit *et al*., 2002). Wings were removed and stored in glassine envelopes as vouchers. Thorax and head were removed and stored in NaCl-saturated dimethyl sulfoxide at –20°C for DNA extraction and processing. Approximately half of the ovaries were dissected immediately, and for the remainder, abdomens were stored in 96% ethanol and transported to the laboratory for fine dissection. In all cases, ovaries were dissected from the abdomen in ice-cold phosphate buffered saline (PBS) using fine forceps and insect pins. Trachaeae and fat bodies were removed manually, and images were taken at 8X, 12.5X, and 20X magnification for phenotyping. Of the ovaries dissected in the field, six backcrosses and two pure *H. pardalinus butleri* were stored in RNALater solution for RNAseq (ThermoFisher AM7020).

### 2.2 Ovary staining and phenotyping

For every dissection, we scored the developmental progress of ovaries on a scale of 0 (empty ovaries) to 3 (containing fully-developed yolky eggs) based on gross morphology (fertility score, Fig. S1, and see examples in Fig. 2). Three images from each ovary were scored blind by two independent scorers. The resulting six scores per ovary were averaged to yield a single fertility score for each individual.

**Figure 2:**
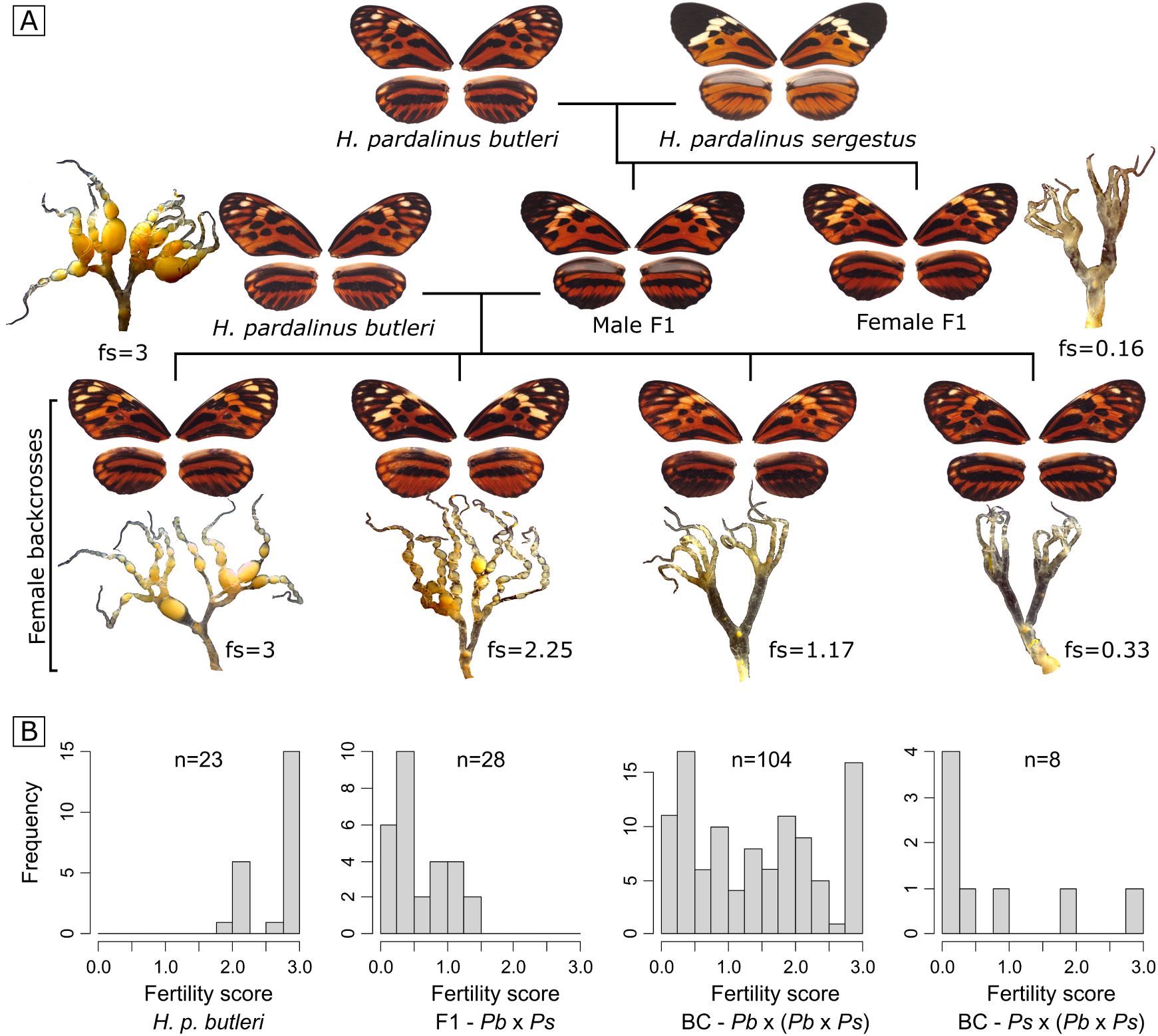
Crossing scheme and distribution of phenotypes. **A** Crossing *H. p. butleri* with *H. p. sergestus* in either direction produces sterile female F1s, while male F1s are fertile. Backcrossing these males in either direction produces females with variable fertility. Example wing phenotypes and dissected ovaries for backcrosses to *H. p. butleri* are shown, with fertile individuals to the left and sterile individuals to the right; fs = fertility score assigned to the dissected ovary. **B** Histograms of ovary fertility scores for i) *H. p. butleri* females, ii) F1s produced by mating a female *H. p. butleri* (*Pb*) with a male *H. p. sergestus* (*Ps*), iii) backcrosses produced by mating fertile male F1 (*Pb* x *Ps*) with female *H. p. butleri* (*Pb*), and backcrosses produced by mating fertile male F1 (*Pb* x *Ps*) with female *H. p. sergestus* (*Ps*).

In a subset of samples, we characterized the earliest arrested developmental stages of oocytes through nuclear staining with DAPI, using the stages described in the silkmoth (*Bombyx mori*) as a reference (Fig. 3). Individual ovarioles from alcohol-stored ovaries were removed and rehydrated by 15 minute incubations in serial dilutions of ethanol in 0.1% tween 20 in 1X phosphate-buffered saline (PBT) (Ethanol concentrations: 95%, 90%, 80%, 60%, 40%, 20%, 0%). Once fully re-hydrated, ovaries were incubated in acridine orange solution (ThermoFisher A1301; 5 µg/mL in PBT) to visualize cytoplasm. They were then washed in PBT before being stained with DAPI (1 µL/mL in PBT), washed once more in PBT, and mounted on slides with VectaShield (Vector Labs). Slides with stained ovarioles were scanned with a Zeiss Axio Scan Z1, and high-magnification images were taken with a Zeiss LSM 880 upright confocal microscope. The most highly developed follicle in each ovariole was staged through visual comparison to oocyte development stages described in *Bombyx mori* (Yamauchi & Yoshitake, 1984).

**Figure 3:**
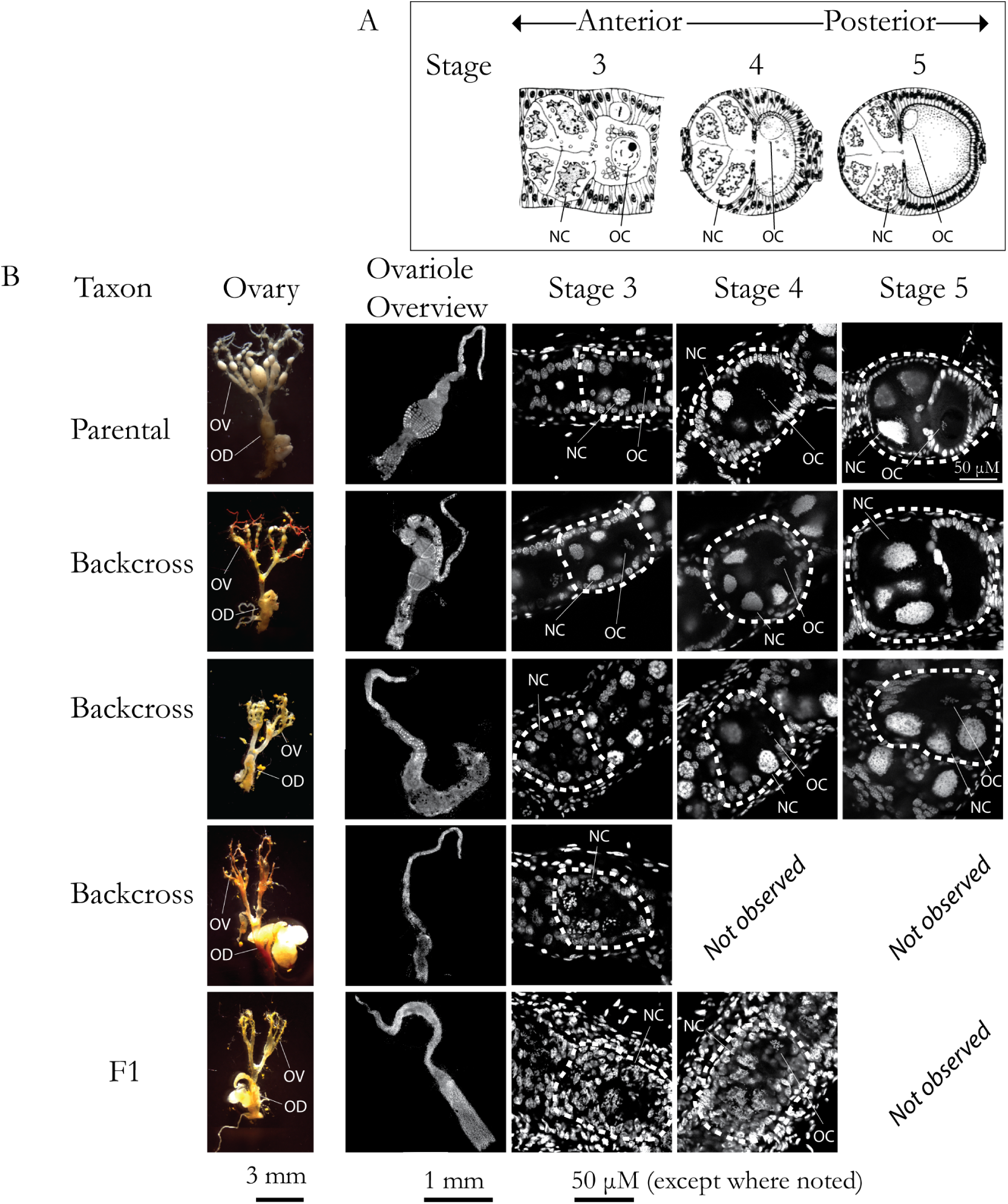
Developing oocytes. **A.** Idealized developing follicle stages (Yamauchi & Yoshitake, 1984) **B.** Brightfield and confocal images of DAPI-stained ovaries. Each row displays an overview image, as well as individual follicles at indicated stages from the same ovary. Scale bars for ovariole overviews are shown below the relevant column. Scale bar for stages 3-5 is shown below the stage 3 column, except where indicated in image. “Not observed” represent stages not present in the illustrated ovariole. In ovary images, one ovariole (OV) and the oviduct (OD) are indicated. Individual follicles are encircled by dashed lines. Where visible, one nurse cell nucleus (NC) and the oocyte cell nucleus (OC) in the highlighted follicle are indicated.

### 2.3 DNA extraction and sequencing

RNA-free genomic DNA was extracted from individuals used in QTL mapping (see below) using a Qiagen DNeasy Blood and Tissue Kit and following the manufacturer’s standard protocol. Restriction site Associated DNA (RADSeq) libraries were prepared using a protocol modified from Etter et al. (Etter *et al*., 2011; Hoffman *et al*., 2014), using a *PstI* restriction enzyme, sixteen 6bp P1 barcodes and eight indexes. DNA was Covaris sheared and gel size selected to 300-700bp. 128 individuals were sequenced per lane, with 125bp paired-end reads, on an Illumina HiSeq 2500.

### 2.4 SNP calling

FastQ files from each RAD library were demultiplexed using process radtags from Stacks (Catchen *et al*., 2013), and BWA-MEM (Li, 2013) was used with default parameters to map the reads both to the *H. melpomene* genome (Hmel2.5) (Davey *et al*., 2017) and to the *H. pardalinus* genome (Hpar) (Seixas *et al*., 2021). BAM files were then sorted and indexed with SAMtools (Li *et al*., 2009), and Picard-tools v 1.119 (https://github.com/broadinstitute/picard) was used to add read groups and mark PCR duplicates. To check for incorrectly labelled samples, we estimated the sex of a sample by dividing the mean number of reads per kilobase on the Z chromosome by the mean value for autosomes. This returned a value close to 1 in males and 0.7 in females, which can then be compared with the recorded sex of the sample. To further check for labelling errors, confirm pedigrees, and assign samples with unrecorded pedigree to families, we used Plink 1.9 (Chang *et al*., 2015) to estimate the fraction of the genome that is identical by descent (IBD; 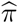) between all pairwise combinations of samples (siblings and parent-offspring comparisons should yield 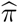 values close to 0.5). In addition, for specimens that were sequenced multiple times in order to improve coverage, we checked that samples were derived from the same individual (with 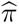 values close to one). We then merged these samples, using the MergeSamFiles command from Picard-tools, and used Samtools’ mpileup command to call single nucleotide polymorphisms (SNPs) for linkage map construction.

### 2.5 Linkage map construction

Linkage maps were built using reads aligned to each of the reference genomes using Lep-MAP3 (Rastas, 2017). The ParentCall2 module was used to correct erroneous or missing parental genotypes, and call sex-linked markers using a log-odds difference of *>*2. We used Filtering2 to remove SNPs showing segregation distortion, specifying a *P* -value limit of 0.01 (i.e., there is a 1:100 chance that a randomly segregating marker is discarded). Because we genotyped only female offspring, we did not filter sex-linked markers for segregation distortion. We then used SeparateChromosomes2 to cluster markers to linkage groups, specifying zero recombination in females and joining pairs of markers with LOD-score greater than 14. To obtain recombination distances between markers, we fixed the order of the markers to their order on the Hmel2.5 or Hpar genome assemblies, and then evaluated this order, again using paternally and dual informative markers. Lep-MAP3 outputs fully informative and phased genotypes with no missing data, which can be used for QTL mapping.

### 2.6 QTL mapping

Genetic data were analysed as backcrosses (Fig. 2) using the paternally inherited allele. We used R/QTL (Broman *et al*., 2003) to estimate genotype probabilities at 1 cM intervals, using the Haldane mapping function and an assumed genotyping error rate of 0.001. Loci with inferred genotypes were labelled using the chromosome and the centimorgan position. We used Haley-Knott (H-K) regression to test for associations between the estimated genotype probabilities at each marker and fertility score (Haley & Knott, 1992). BB genotypes were coded as 0.5 and BS genotypes were coded as -0.5, where B is the *H. p. butleri* allele and S is the *H. p. sergestus* allele.

We first built a single locus additive QTL model at each position in the genome (H_1_; *y = µ*_1_ + *β*_1_*q*_1_ + *ε*) and calculated the log_10_ likelihood ratio (LOD score) comparing (H_1_) with the null hypothesis of no QTL (H_0_; *y = µ*_1_ + *ε*). To identify loci that act in combination to produce the phenotype, we then estimated LOD scores using all pairwise combinations of typed markers and inferred genotypes at 1 cM intervals across the genome, while allowing interactions between them (H*_f_; y = µ*_1_ + *β*_1_*q*_1_ + *β*_2_*q*_2_ + *β*_3_*q*_1_*q*_2_ + *ε*). The difference between LOD values for (H*_f_*) and the corresponding two locus additive model (H*_a_; y = µ*_1_ + *β*_1_*q*_1_ + *β*_2_*q*_2_ + *ε*) gives the improvement in fit attributable purely to interactive effects (H*_int_*). The difference between LOD*_f_* and the maximum LOD value obtained from single QTL locus models at either marker indicates the presence of a second QTL, allowing for epistasis (H*_fv_*_1_). We also performed these analyses while controlling for kinship. To do this, we used LepMap to estimate 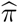 (IBD) between all individuals. We then created a variance-covariance matrix of genetic relatedness, and included this in our models as a random effect. Significance of QTL scans was assessed by permuting the phenotypes relative to the genotypes (10,000 permutations). Because we analysed only female backcrosses, the degrees of freedom for QTL models at sex-linked and autosomal loci are the same, and so we set a single genome-wide significance threshold for each scan.

### 2.7 Population genomics

To examine genomic differentiation between the *H. p. sergestus*, *H. p. butleri* and *H. elevatus*, previously published whole genome re-sequencing data (four individuals each taxon) were used (NCBI accession numbers: ERS070236; ERS977673; ERS977674; ERS070238; ERS4368504; SRS329822; SRS329823; SRS329824; SRS329825; SRS329826; SRS3298233; SRS1247739; ERS235668; ERS977715; ERS977716; ERS977717). Raw reads were filtered for Illumina adapters using cutadapt (Martin, 2011) and mapped to the Hmel2.5 (Davey *et al*., 2017) (Seixas *et al*., 2021) genomes using BWA MEM v0.7.15. Duplicate reads were removed using sambamba v0.6.8 (Tarasov *et al*., 2015) and the Genome Analysis Toolkit (GATK) v3.8 RealignerTargetCreator and IndelRealigner modules (DePristo *et al*., 2011; McKenna *et al*., 2010) were used to realign reads around indels. Genotype calling was performed for each taxon separately with bcftools (Li *et al*., 2009) mpileup and call modules (Li, 2011), using the multiallelic and rare-variant calling option (-m) and requiring a minimum mapping quality and base quality of 20. Genotype calls were required to have a minimum quality score (QUAL) of 20, RMSMappingQuality (MQ) *≥* 20, genotype quality (GQ) *≥* 20 and a minimum individual depth of coverage (DP) *≥* 8 (or DP *≥* 4 for the Z chromosome of females). Genotypes within 5 bp of an indel were recorded as missing data.

Differentiation (*F_ST_*), pairwise genetic distances (*D_XY_*) and nucleotide diversity(*π*) between the three taxa studied were estimated along the genome in overlapping 25 kb windows (with 5 kb steps) using the popgenWindows.py script (available from https://github.com/simonhmartin/genomics_general).

### 2.8 RNA extraction and sequencing

Ovaries stored in RNALater were further dissected into pre-vitellogenic (i.e., before yolk deposition) follicles, vitellogenic follicles, and choriogenic follicles + chorionated eggs (the chorion is the proteinaceous “eggshell” of an insect egg). Each of these three subsets was processed separately. Tissue was blotted dry with Kimwipes to remove excess RNALater solution, transferred to TRIZOL and homogenized with the PRO200 tissue homogenizer (PRO Scientific). RNA was extracted with the Direct-zol RNA miniprep kit (Zymo R2051). The mRNA libraries were prepared by the Harvard University Bauer Core with the KAPA mRNA HyperPrep kit, with mean fragment insert sizes of 200-300bp, and were sequenced on a NovaSeq S2, producing an average of 49 million paired-end, 50 bp reads per library (Table S2).

RNASeq reads were mapped to the *H. melpomene* v2.5 transcriptome (Pinharanda *et al*., 2019) using kallisto (Bray *et al*., 2016). Approximately 70% of reads were mapped to the transcriptome per sample, and that value did not differ between the *H. pardalinus butleri* samples and the backcrosses (Table S2). Aligned reads were normalized to account for sequencing coverage, transcript length, and RNA composition using sleuth (Pimentel *et al*., 2017). Raw counts were log-transformed, and expression differences were calculated by comparing the likelihood of the model: *ln*(*counts*) *∼* 1 to the model *ln*(*counts*) *∼*1 + *binaryscore* (Pimentel *et al*., 2017).

In order to identify conserved genes expressed in butterfly oogenesis, we used BLAST to identify *H. melpomene* transcripts orthologous to genes expressed in the ovarian transcriptome of the Speckled Wood butterfly *Parage aegeria* (Carter *et al*., 2013). In addition, we used OrthoFinder (Emms & Kelly, 2019) to identify transcripts with orthologous genes in *Drosophila melanogaster*, and then filtered this list with the keywords “oogenesis” OR “follicle” OR “nurse” OR “oocyte” using the phenotypic data on Flybase (http://flybase.org).

## 3 Results

We reared 143 F1 hybrid offspring of *H. p. butleri* and *H. p. sergestus*. Female F1s in both directions of cross were sterile. To investigate the molecular and genetic basis of hybrid sterility between the two populations, we backcrossed fertile F1 hybrid males to both parental species, rearing 320 offspring. F1 and backcross broods eclosed with approximately equal sex ratios (69 females:73 males and 164 females:156 males, respectively), which suggests a lack of sex bias in immature stage viability.

Almost all individuals from parental populations contained developing follicles that reached the final stages of vitellogenesis, and most had fully developed eggs (*n*=11/12). However, ovaries of F1 hybrids seemed devoid of developing oocytes (Fig. 2). Female backcrosses with *H. p. butleri* mothers yielded an approximately bimodal distribution of ovary phenotypes (Fig. 2B), while a small samples of backcross females (n=8) to *H. p. sergestus* exhibited a skewed distribution, with mostly sterile individuals (Fig. 2B).

All F1 and backcross individuals had early-stage follicles, but sterile individuals showed arrested development after oocytes reached approximately stage 3. This stage marks a developmental timepoint after oocyte vs. nurse cell differentiation and follicle formation, but before vitellogenesis (yolk deposition) (Yamauchi & Yoshitake, 1984; Büning, 1994). Using the 42 individuals for which we could confidently assign the latest developmental stage and fertility score, we verified that the two metrics were highly correlated (logistic regression, *p* = 2.45 *×* 10*^−^*^11^, Supplementary Fig. S2).

### 3.1 QTL mapping

We sequenced 87 females from 7 families produced by backcrossing F1 males to *H. p. butleri* females. Using RADSeq reads aligned to Hmel2.5 reference genome, the linkage map for these individuals comprised 124,456 markers across 21 chromosomes, with a total map length of 1106.95 cM (Supplementary table S1 and Figs. S3-S6).

Scanning the genome for additive, single locus QTLs associated with fertility score (H_1_) revealed a broad central region on the Z chromosome (Figs. 4, S8, Tables 1, S3). The maximum LOD value was observed at 29.2 cM (Fig. 4B, C), with mean predicted fertility scores of 1.81 for the *H. p. butleri* allele and 0.93 for the *H. p. sergestus* allele (*R*^2^ = 0.20). The *H. p. butleri* allele had higher predicted fertility scores than the *H. p. sergestus* allele all along the Z chromosome, but the difference declined to nearly zero towards the distal end of the chromosome.

**Figure 4:**
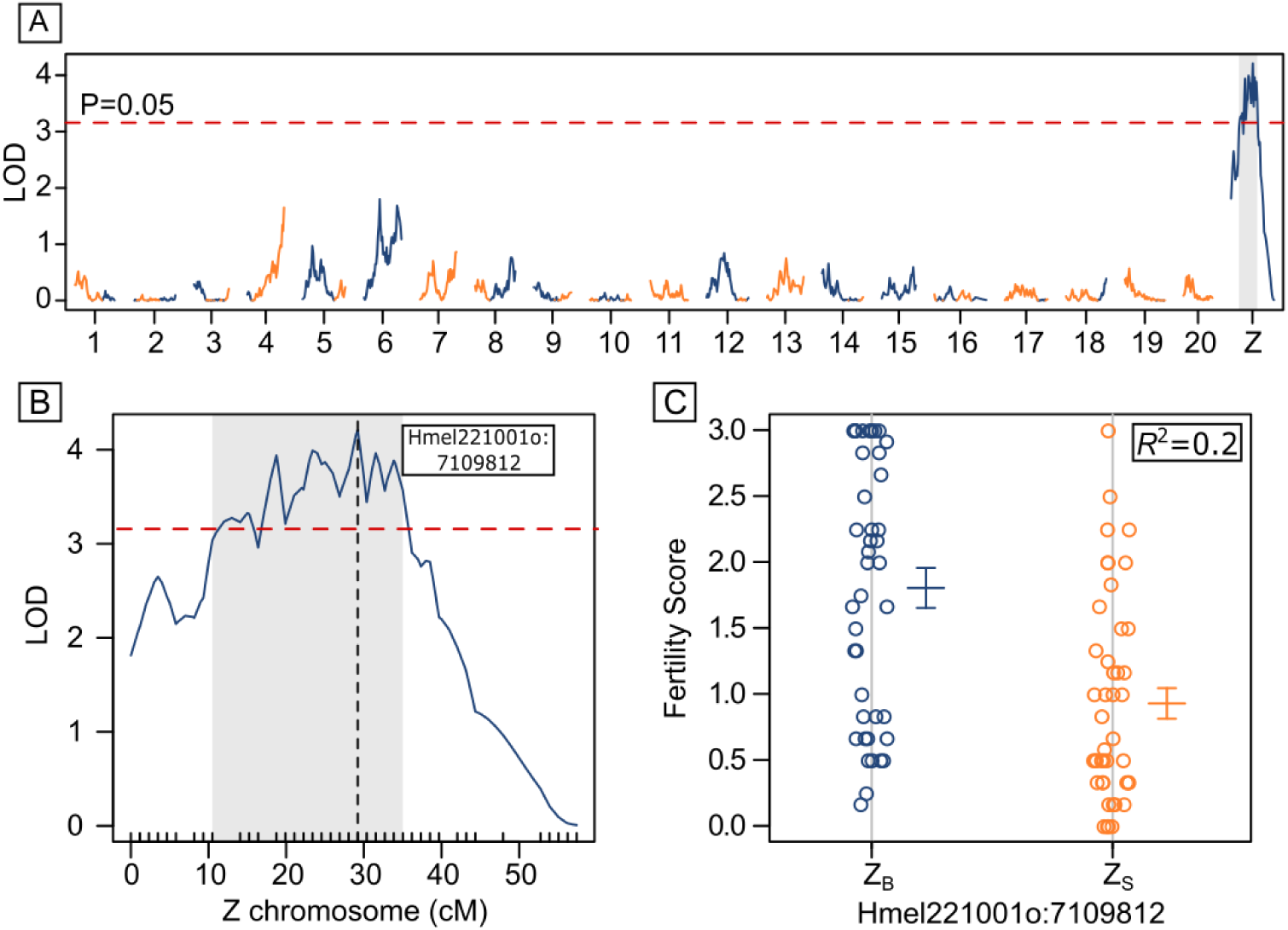
Single QTL analysis. **A.** LOD values at each marker across the genome, calculated using H-K regression and with reads aligned to Hmel2.5. The red dashed line indicates the genome-wide significance threshold (p*<*0.05; 10,000 permutations), and the grey shaded area the Bayesian credible intervals for the peak on Z. Lines are coloured depending on whether the *H. p. butleri* allele (blue) or the *H. p. sergestus* allele (yellow) had higher fertility. **B.** Enlargement of Z chromosome, with the QTL peak at 29.21 cM indicated by the vertical dashed line (corresponding to physical position Hmel221001o:7109812). **C.** Fertility scores at the QTL peak Z markers are hemizygous and coded by a single letter (B = *H. p. butleri* and S = *H. p. sergestus*), and explain 20% of the variance in fertility score. Errors bars are standard errors.

**Table 1:** Summary of significant single locus (H_1_) and two locus (H_f_) QTL models (using reads aligned to Hmel2.5). The first column gives the LOD value of the full model (H*_f_*), with the significance estimated by permutation (+P*<*0.1, *P*<*0.05, **P*<*0.01, ***P*<*0.001). The next columns are the chromosome and QTL marker (scaffold and median physical position within the peak). The centimorgan limits are the Bayesian credible intervals, and the physical limits are the nearest typed flanking markers of that interval (all physical limits were on the same scaffold as the QTL peak, except for the chromosome 4 interaction † with Z, which was on scaffold Hmel204003 of Hmel2.5). The final five columns give the parameter estimates and *R*^2^ of the model. *β*_1_*q*_1_ and *β*_2_*q*_2_ are the estimated additive effects for the QTLs, i.e. the difference between the average fertility scores for the alternative genotypes, and *β*_3_*q*_1_*q*_2_ is the coefficient for the interaction between the 2 loci. Model coefficients comprise the estimated value, the standard error, and the significance (thresholds as above). The significant interaction between chromosome 8 and 20 was detected using individuals holding a Z*_SB_* chromosome only.

| $LOD_f$ | chr | QTL1<br>marker | cM | limits<br>(cM) | limits<br>(physical) | chr | QTL2<br>marker | cM | limits<br>(cM) | limits<br>(physical) | $\mu_1$ | $\beta_1 q_1$ | $\beta_2 q_2$ | $\beta_3 q_1 q_2$ | $R^2$ |
| --- | --- | --- | --- | --- | --- | --- | --- | --- | --- | --- | --- | --- | --- | --- | --- |
| 4.21** | Z | Hmel221001o:<br>7109812 | 29.21 | 10.49-35.02 | 4710269-7966755 |  |  |  |  |  | 1.370.09 | 0.88±<br>0.19*** |  |  | 0.2 |
| 6.72+ | 4 | Hmel204001o:<br>9208991 | 49.03 | 25.72-49.03 | 6102423-7036† | Z | Hmel221001o:<br>7109812 | 29.21 | 15-36.19 | 5201659-7976401 | 1.32±<br>0.09*** | -0.56±<br>0.18** | 0.81±<br>0.18*** | -0.59±<br>0.37 | 0.3 |
| 6.65+ | 12 | c12.loc25 | 25 | 17.47-55.93 | 5100627-16319705 | Z | Hmel221001o:<br>5752071 | 18.75 | 3.49-33.86 | 2210592-7862353 | 1.39±<br>0.09*** | 0.44±<br>0.18* | 0.87±<br>0.18*** | 1±<br>0.36** | 0.3 |
| 7.87** | 15 | Hmel215003o:<br>8041294 | 38.48 | 10-44.3 | 4230382-9907975 | Z | Hmel221001o:<br>5752071 | 18.75 | 14.07-30 | 4795563-7180089 | 1.34±<br>0.09*** | 0.29±<br>0.18 | 0.7±<br>0.18*** | 1.47±<br>0.36*** | 0.34 |
| 14.52*** | Z | Hmel221001o:<br>3045330 | 4.65 | 2.33-5.81 | 2179644-4463341 | Z | Hmel221001o:<br>10565964 | 55.08 | 53.91-56.24 | 8861371-13311117 | 1.4±<br>0.07*** | 0.6±<br>0.15*** | -0.03±<br>0.15 | -2.53±<br>0.3*** | 0.54 |
| 7.76* | 8 | Hmel208001o:<br>1005579 | 9.3 | 8.14-10 | 618400-1390337 | 20 | Hmel220003o:<br>5817143 | 11.86 | 6-12 | 3012264-6466723 | 1.61±<br>0.08*** | 0.24±<br>0.16 | -0.22±<br>0.16 | 2.68±<br>0.32*** | 0.79 |

When scanning for interacting QTLs we identified a negative interaction between a pair of loci at opposing ends (*∼*5 cM and *∼*55 cM) of the sex chromosome, with the full epistatic model explaining 54% of the variance in fertility score (Fig. 5, Tables 1, S3). This pair of loci was highly significant (P*<*0.001) irrespective of whether we tested the combined additive effects and interaction (H*_f_*), the additive effect of the second locus plus the interaction (H*_fv_*_1_) or the interaction alone (H*_int_*), and was robust to family-specific effects (Fig. S8). Recombinant Z chromosomes (Z*_BS_* or Z*_SB_*) had higher fitness (i.e., greater average fertility scores) than either non-recombinant, (Z*_BB_* or Z*_SS_*) (Fig. 5). In addition to the interacting loci on the sex chromosome, we further identified significant pairs of QTLs between the Z chromosome and chromosomes 4, 12 and 15 (Table 1, Fig. 5). We then tested for the single QTL at 29.2 cM on the sex chromosome while controlling for the epistatically interacting pair of QTLs at either end. It remained significant, but its position shifted slightly to 33.86 cM. Bringing these three QTLs together in a single model (y = *µ*_1_ + *β*_1_*q*_1_ + *β*_2_*q*_2_ + *β*_3_*q*_3_ + *β*_4_*q*_1_*q*_2_ + *ε*) explained 62% of the variance in fertility score.

**Figure 5:**
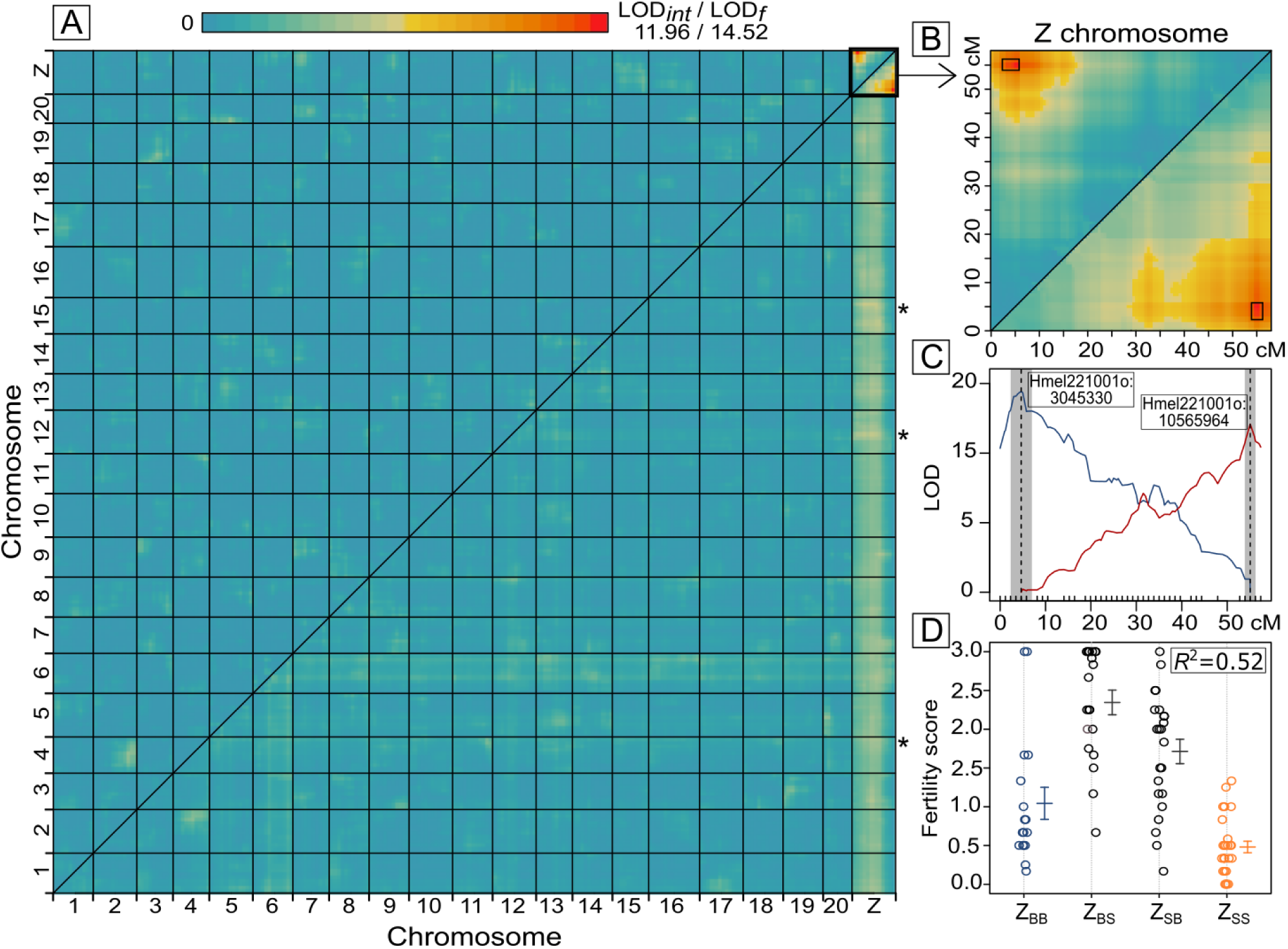
Multiple QTL analysis. **A.** Heat map for LOD*_f_* values (the full model; lower right triangle) and LOD*_int_* scores (the interaction component; upper left triangle) between pairwise combinations of markers across the genome, using H-K regression and reads aligned to Hmel2.5. Blues indicate low scores, reds indicate high scores (maximum observed LOD*_int_* = 11.96, maximum observed LOD*_f_* = 14.52). Statistically significant LOD*_f_* values between the Z chromosome and the autosomes are marked with an asterisk. **B.** Enlargement of the Z chromosome, with the Bayesian credible intervals of the significant interaction shown as black boxes. **C.** Profile LOD curves for the epistatic QTL on Z chromosome, with the blue line for the proximal QTL and the red line for the distal QTL. The vertical dotted lines give the positions of the QTL peaks, and the grey shaded errors indicate the Bayesian credible intervals. The physical positions of the markers at the QTL peaks are shown in the text boxes. **D.** Fertility scores for 87 backcross individuals grouped by their haplotypes at the two interacting markers on the Z chromosome (Hmel221001o:3045330 and Hmel221001:10565964). These four haplotypes explain 52% of the variance in fertility score. Unrecombined pairs of markers inherited from *H. p. butleri* (Z*_BB_*) or *H. p. sergestus* (Z*_SS_*) are coloured blue and orange, respectively. Errors bars are standard errors.

To understand these results further, we divided individuals into four groups depending on their genotypes at the two interacting loci on the Z chromosome (Z*_BB_*, Z*_SS_*, Z*_BS_*, Z*_SB_*). For each of these groups, we then plotted fertility against the fraction of the autosomes homozygous for *H. p. butleri* alleles (*B/B*). We hypothesised that if sterility is driven by interactions between the Z chromosome and autosomes, this fraction should be positively correlated with fertility score for those individuals holding a *Z_BB_*. As expected, for *Z_BB_* individuals, we found a significant positive correlation between the proportion of autosomal markers derived from *H. p. butleri* (Fig. 6A). Interestingly, we also found a significant negative correlation for Z*_SB_* individuals. We then conducted QTL mapping on each of these groups. For individuals with a recombinant Z*_SB_* chromosome, we identified a significant interaction (LOD*_int_* = 6.97, P*<*0.01, *R*^2^ = 0.79) between loci at 9.3 cM on chromosome 8 and 11.9 cM on chromosome 20 (Fig. 6B-D). No significant QTLs were detected for the other subgroups (Z*_BB_*, Z*_SS_* and Z*_BS_*).

**Figure 6:**
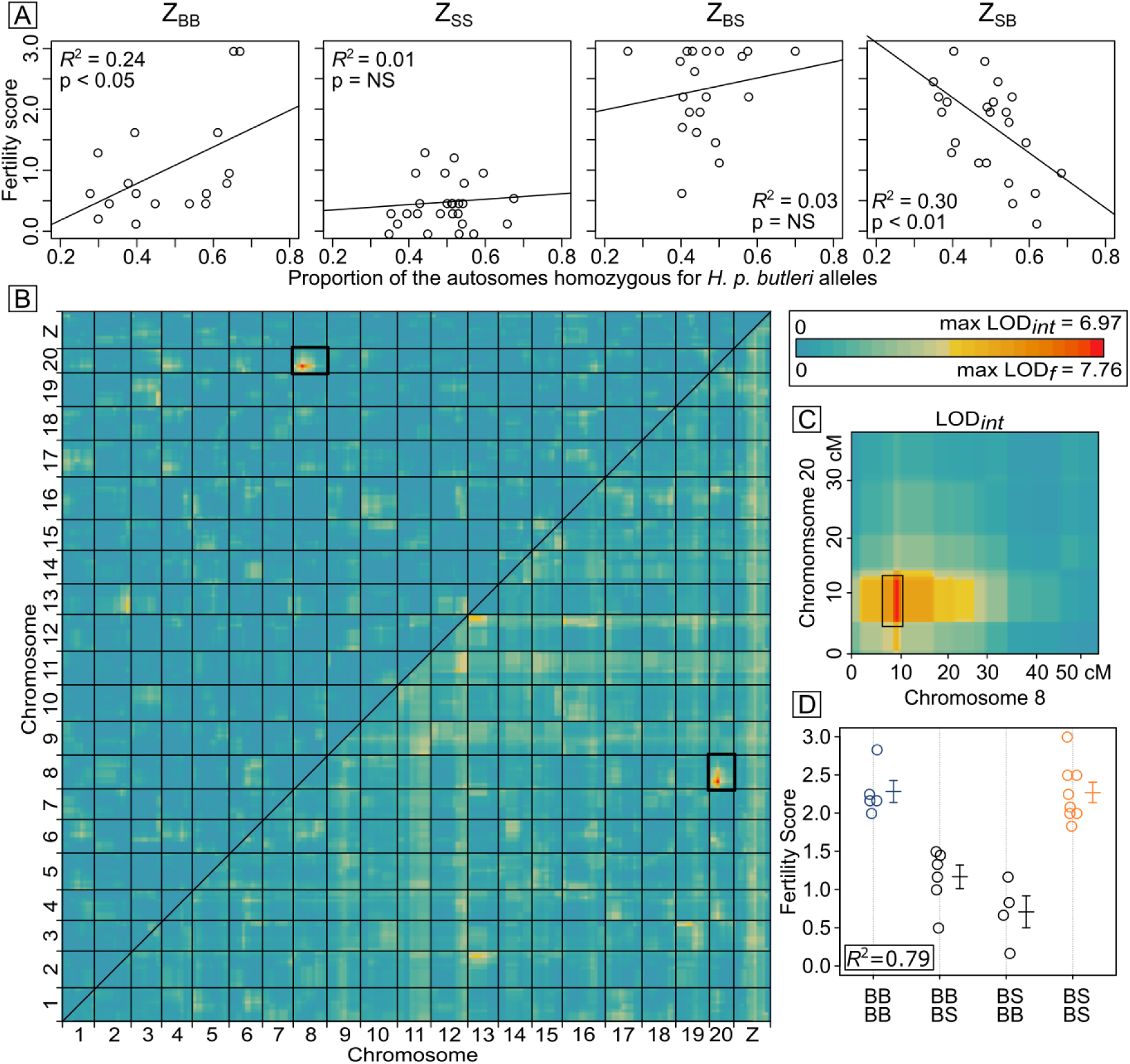
Analysis of Z linked epistatic markers. **A.** For each Z chromosome haplotype (Z*_BB_*, Z*_SS_*, Z*_BS_*, Z*_SB_*), the proportion of the autosome that is homozygous for *H. p. butleri* alleles was plotted against fertility score. **B.** Heat map for two dimensional QTL scan using only Z*_SB_* individuals. LOD*_f_* values are shown in the lower right triangle) and LOD*_int_* values in the upper left triangle. The highlighted box shows the significant associations identified between chromosomes 8 and 20. **C.** Enlargement of LOD*_int_* between chromosome 8 and chromosome 20, with the Bayesian credible intervals of the QTLs shown as black boxes. **D.** Fertility scores for the four autosomal genotypes of Z*_SB_* individuals, with the genotype at chromosome 8 (Hmel208001o:1005579) written above, and the genotype at chromosome 20 (Hmel220003o:5817143) written below. These genotypes explain 79% of the variance in fertility score. Errors bars are standard errors.

### 3.2 Population genomics of the Z chromosome

Nucleotide diversity (*π*) in *H. p. sergestus* was low along the most of the Z chromosome, but higher in *H. p. butleri* and *H. elevatus*, which were near identical (Fig. 7A). Pairwise genetic differentiation (*D*_xy_) was very similar between all three taxa, barring a 250 kb region in the center of the Z chromosome (6.5-6.75 MB) between *H. p. butleri* and *H. elevatus*, where it dropped close to zero (Fig. 7B). This region was also characterised by high *F*_ST_ between *H. p. sergestus* and *H. p. butleri*, which falls in the centre of the additive QTL peak (Fig. 7C). *F*_ST_ was generally elevated at the ends of the Z chromosome as well, possibly due to the two epistatic QTLs; however, these regions also have low rates of recombination (Fig. 7C), which can lead to high *F*_ST_ values even in the absence of selection (Burri, 2017). Overall, the mean *F*_ST_ between *H. p. butleri* and *H. p. sergestus* for the Z chromosome was 0.37, making it the most divergent chromosome. The autosomes ranged from 0.23 (chromosome 3) to 0.35 (chromosome 19), with an overall mean of 0.27.

**Figure 7:**
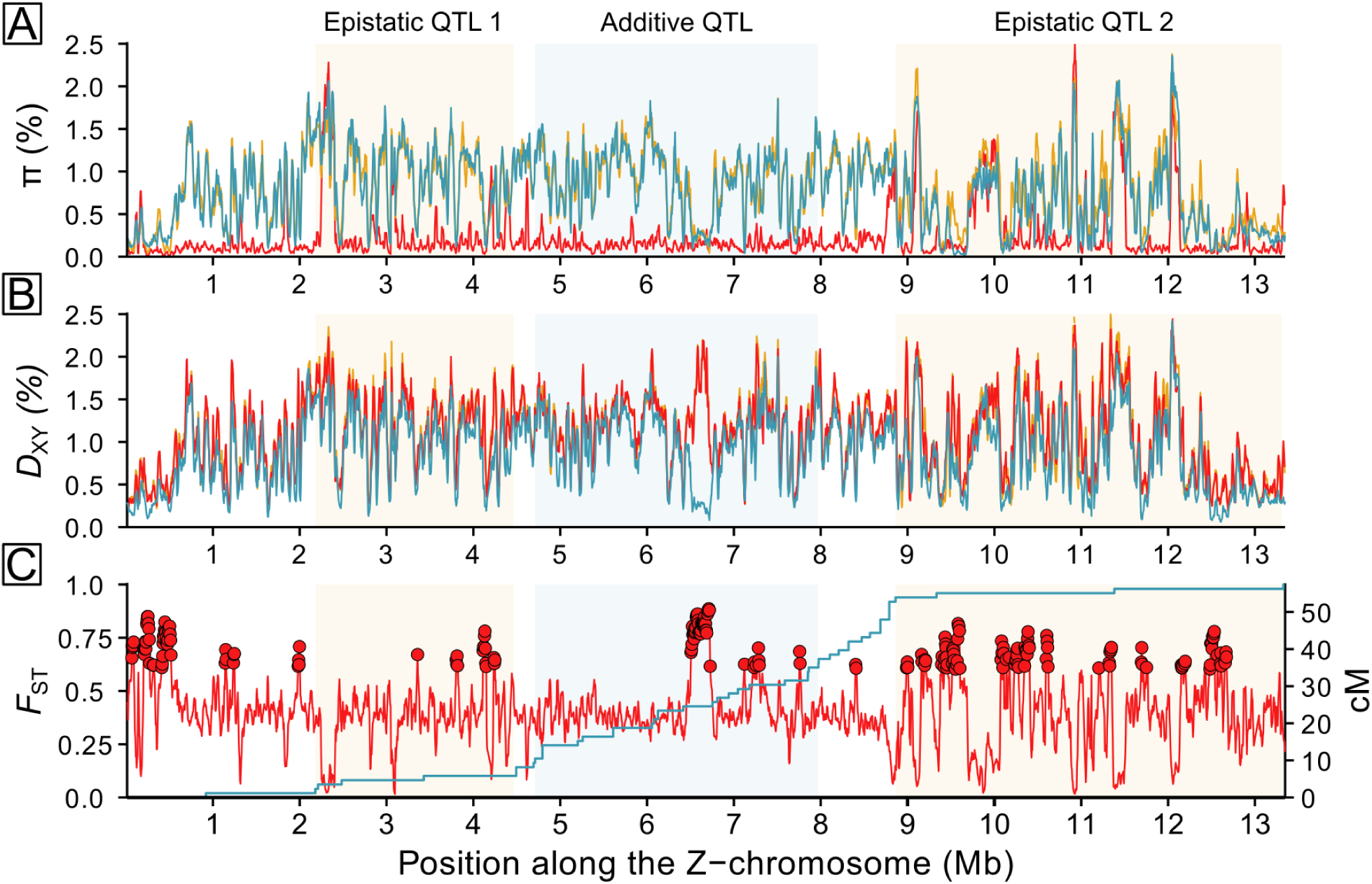
Population genetic summary statistics and recombination rate along the Z chromosome. **A.** Nucleotide diversity (*π*) within *H. p. sergestus* (red), *H. p. butleri* (yellow), and *H. elevatus* (blue). **B.** Mean pairwise absolute genetic distance (*D*_xy_) between *H. p. butleri* and *H. p. sergestus* (red), *H. p. butleri* and *H. elevatus* (blue), and *H. p. sergestus* and *H. elevatus* (yellow). **C.** Genetic differentiation (*F*_ST_; red line) between *H. p. butleri* and *H. p. sergestus*, with genome-wide *F*_ST_ outliers as points, based on Z-scores *>* 3. The blue line shows genetic distance (cM) plotted against physical distance (Mb). Shaded areas correspond to the Bayesian credible intervals for the two epistatic QTL at 4.65 and 55 cM, and the single additive QTL at 29.21 cM. *D*_xy_ and *F*_ST_ were calculated in sliding windows of 25 kb (with 5 kb increments).

### 3.3 Differential expression analysis

The dysgenic sterility phenotype is first evident in early stage, pre-vitellogenic oocytes (Fig. 3). We focused on this region of ovaries in quantifying RNA expression differences among backcross individuals. We dissected the pre-vitellogenic (approx. stage 3 and earlier) follicles from six backcross ovaries, two of which were assigned a fertility score of 0-1, two of 1-2, and two of 2-3. We micro-dissected pre-vitellogenic follicles to investigate the specific phenotype of developmental failure in early-stage oocytes, and further classified the phenotypes with a binary scheme: 0 for absence of vitellogenic follicles, 1 for presence of any vitellogenic follicles. In each case, tissue was dissected from all four ovarioles of a single ovary, and we acquired approximately 49 million reads per individual. After filtering our data for sequencing and mapping quality, we quantified expression of 16,774 transcripts (Fig. 8).

**Figure 8:**
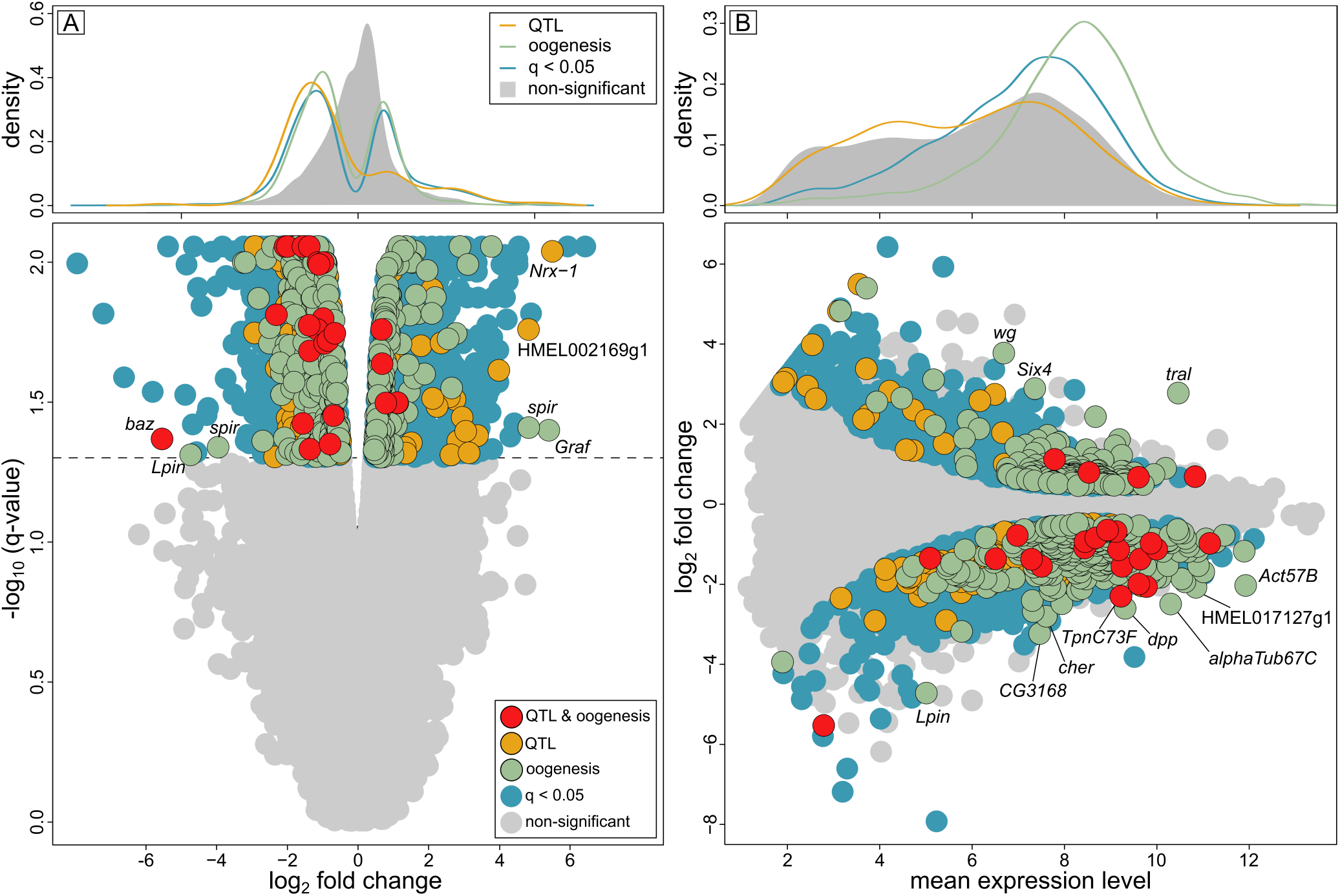
Backcross differential expression. **A.** Volcano plot. The change in expression between fertile (fertility score *>* 1.5) and sterile (fertility score *≤* 1.5) ovaries is plotted against the q-value. Positive values of fold change imply higher expression in fertile ovaries. Significantly differentially expressed transcripts are shown in blue, of which those with orthologues implicated in oogenesis in either *D. melanogaster* or *P. aegeria* are in green, those within the Bayesian credible intervals of QTLs on chromosomes 8, 20 and Z are shown in orange, and those fitting all these criteria are in red. QTL and/or oogenesis outliers (those with absolute fold change values in the top 1% of all transcripts) are labelled. Non-sigificant transcripts are in grey. The density plot shows the distribution of fold change values i) implicated in oogenesis, ii) found within QTLs, and iii) significantly differentially expressed in fertile/sterile individuals, with the density of all transcripts shaded in grey. Interestingly, most QTL transcripts are overexpressed in the sterile ovaries. **B.** Mean expression of transcripts plotted against fold change in expression. Symbols as in A, except the density plot shows the distribution of mean expression levels. Labelled QTL and/or oogenesis outliers were defined as those falling in the top 1% of transcripts ranked using mean expression *×* fold change.

We then carried out a principal components analysis of these expression data. PC1, which explains over 50% of variance, separates the three fertility score categories in order (Fig. 8B). We performed a Wald test to evaluate the effect of change in expression of each transcript to the fertility phenotype in the backcrosses (Chen *et al*., 2011). After correcting for multiple comparisons, a total of 14%, or 2315 transcripts showed significant effects of expression on binary phenotype (*q <* 0.05) (Fig. 5C,D). Of these, 941 displayed a positive association with development, meaning that the transcript was expressed at a higher level in more highly developed ovaries. The remaining 1386 differentially expressed transcripts displayed a negative association with development. To narrow our list of candidate genes, we filtered our differentially expressed transcripts for genes implicated in butterfly oogenesis. We identified 1,771 transcripts in the *H. melpomene* transcriptome that gave strong BLAST hits to genes expressed in *Pararge aegeria* eggs and ovaries (Carter *et al*., 2013). As expected, these genes showed generally high expression levels in the sampled transcriptomes relative to other genes (Fig. 8A,B). 306 (17%) of these genes were also differentially expressed in backcrosses with different developmental phenotypes. One of the transcripts, *Trailer hitch* (*tral*), has high overall expression, strong differential expression, and is known to be involved in oogenesis of *D. melanogster* and *P. aegeria* (Wilhelm *et al*., 2005; Carter *et al*., 2013) (Fig. 8).

We then searched within the Bayesian credible intervals of the QTLs for differentially expressed transcripts with orthologs implicated in oogenesis in either *D. melanogaster* or *P. aegeria* (Fig. 8). Applying this approach to the two interacting QTLs on the Z chromosome, we identified one candidate gene (*magu*) in the first QTL 4.65 cM, and eight in the second QTL at 55 cM (*Egfr*, *fax*,*Gs2*, *Nedd8*, *parvin*, *Prm*, *sls*, *Syx7*). Within the central additive QTL on the Z chromosome at 29.2 cM, we found two candidate genes (*trol* and *csw*). In the highly region divergent region within this QTL (6.5 - 6.75 Mb, Fig. 7) there are 14 genes, one of which has an orthologue (*ncd*) required for spindle assembly in oocytes in *Dropsophila* (Endow & Komma, 1997). However only three were significantly differentially expressed among fertile and infertile hybrids, and none of those had orthologs implicated in oogenesis.

Within the QTL at 11.86 cM on chromosome 20, we identified 11 candidates (*baz*, *CG12104*, *CG1572*, *CrebB*, *Ect4*, *Eip75B*, *ine*, *mys*, *Pitslre*, *Ran*, *TpnC73F*). In the QTL at 9.3 cM on chromosome 8, there were only 3 differentially expressed transcripts, only one of which had an orthologue known to be involved in oogenesis (*Art1*). However, one transcript (HMEL037834g1.t2, with the orthologue *Nrx-1*) stood out due to very high fold change (*β* = 5.51) between sterile and fertile individuals, and its physical position (997,675 - 1,074,578 Mb) falls in the centre of the peak of the QTL (844,849 - 1,232,231 Mb). Genes involved in oogenesis and those significantly differentially expressed between ovaries of varying development were skewed towards being overexpressed in ovaries of females with low fertility scores. This pattern was even more extreme among all genes in QTL intervals, regardless of their function (Fig. 8A). This could mean that the high expression is due to a general phenomenon such as increased chromatin availability, or derepression of transcriptional regulators.

## 4 Discussion

Here, we show that crossing *H. p. butleri* and *H. p. sergestus* in both directions results in F1 hybrid females that are sterile due to disrupted oocyte development, and QTL analysis of backcrosses to *H. p. butleri* shows that sterility is sex-linked. We identify a strong epistatic interaction between loci at opposite ends of the Z chromosome, and a broader, additive QTL towards the centre. In addition, we identify an epistatic interaction involving the Z chromosome and chromosomes 8 and 20, as well as significant associations linking the Z chromosome with chromosomes 4, 12 and 15. By intersecting these with the results of differential expression analysis, we identify a number of candidate genes.

### 4.1 Genetics of hybrid incompatibility in *Heliconius pardalinus*

To our knowledge, this is the first study of Haldane’s Rule in Lepidoptera using modern genomic techniques to demonstrate a complex, epistatic basis of hybrid sterility, as predicted in the Dobzhansky-Muller model. Hybrids between *H. p. butleri* and *H. p. sergestus* are also consistent with the “two rules of speciation” (Coyne & Orr, 1989b). The first of these is Haldane’s rule - the tendency for greater sterility/inviability in the heterogametic sex than in the homogametic sex. There is general consensus that Haldane’s rule can be explained in part by dominance theory, which proposes that interactions between recessive X- or Z-linked alleles from one species and a hybrid autosomal genetic background cause incompatibilities in the heterogametic sex (Coyne & Orr, 2004). Although our data are consistent with dominance theory, other processes, such as faster evolution of Z-linked genes (Charlesworth *et al*., 1987; 2018), may also have played a role in the evolution of hybrid sterility. However, because sterility manifests in females, the “faster male” hypothesis (for example due to sexual selection) can be ruled out in Lepidoptera (Orr & Turelli, 1996; Wu & Davis, 1993).

The second rule of speciation is the “large X effect” on hybrid incompatibility (in Lepidoptera, this is a large effect of the Z chromosome). In hybrids between *Drosophila mauritiana* and *D. sechellia*, the X chromosome has about four times more hybrid male sterility factors than a comparably sized autosomal region (Masly & Presgraves, 2007), and X-linked loci are involved in female, as well as male, hybrid sterility in the *D. virilis* group (Orr & Coyne, 1989). There is, in addition, a large X-effect in taxa with undifferentiated sex chromosomes (Dufresnes *et al*., 2016; Hu & Filatov, 2016), and a large Z effect in birds (Ellegren, 2009). In Lepidoptera, sex-linked hybrid sterility has been shown in *Colias* and *Heliconius* (Grula & Taylor Jr, 1980; Jiggins *et al*., 2001; Naisbit *et al*., 2002), and in general the Z chromosome appears to be a hotspot for genetic differences between species (Prowell Pashley, 1998; Sperling, 1994). Here we document three sex-linked QTLs, suggesting a large effect of the Z chromosome on hybrid sterility in *H. pardalinus* (but see Coyne & Orr (1989b) and Hollocher & Wu (1996) for caveats). The Z chromosome also has a highest mean *F*_ST_ of any chromosome (1.45 times greater than the mean across all the autosomes), consistent with other population genomic studies of butterfly and bird species (Backström & Väli, 2011; Van Belleghem *et al*., 2018). Although higher *F*_ST_ on the Z chromosome is expected from its lower effective population size (Presgraves, 2018), in combination with the sex-linked QTL it is consistent with greater selection against introgressed Z-linked hybrid incompatibilities than on autosomes.

In *Heliconius melpomene*, crosses between Guiana and Central American populations show hybrid female sterility in only one direction of cross (Jiggins *et al*., 2001). This kind of asymmetry in hybrid sterility is expected when Dobzhansky-Muller incompatibilities are relatively few, due to recent divergence (Muller, 1942; Turelli & Moyle, 2007). In *H. pardalinus*, crosses in both directions between *H. p. sergestus* and *H. p. butleri* produce sterile hybrid females, suggesting a more complex, multilocus cause of hybrid sterility. Moreover, if hybrid female sterility arises due to epistatic interactions between the Z chromosome and autosomes, there must be autosomal loci at which *H. p. butleri* alleles are dominant, and others at which *H. p. sergestus* alleles are dominant.

The observation that individuals with unrecombined Z chromosomes (Z*_BB_* and Z*_SS_*) have lower average fertility than recombined Z chromosomes (Z*_BS_* and Z*_SB_*) supports this (Fig. 5D). A Z chromosome inherited from *H. p. sergestus* (Z*_SS_*) will have deleterious interactions with any autosomal loci where *H. p. butleri* alleles are dominant, and so in a backcross to *H. p. butleri*, individuals carrying such a chromosome should never have full fitness. Similarly, individuals with a Z chromosome inherited from *H. p. butleri* (Z*_BB_*) should also have reduced fertility, because of deleterious interactions with autosomal loci with a dominant *H. p. sergestus* allele. However, in a backcross *H. p. butleri*, some fraction of offspring bearing unrecombined *H. p. butleri* Z chromosomes should be fully fertile; those that happen to be homozygous for *H. p. butleri* alleles at all *H. p. sergestus* autosomal dominant loci that interact with the Z. Indeed, two individuals do; these are clearly visible as outliers in Fig. 5D and as predicted they have the highest proportion of their autosomes homozygous for *H. p. butleri* alleles (*B/B*) (Fig. 6A).

It is less easy to explain why individuals holding a recombined Z*_BS_* or Z*_SB_* chromosome on average have higher fertility than uncrecombined chromosomes. Male hybrid sterility between Bogotá and US populations of *Drosophila pseudoobscura* is the product of complex epistasis between seven genes which includes interactions between sex linked markers (Orr & Irving, 2001; Phadnis, 2011). Subsequent work on *D. pseudoobscura* and *D. persimilis* has shown that espistasis can even modify the dominance of loci causing hybrid male sterility (Chang & Noor, 2010). Given this potential for complexity, a complete explanation of the epistatic interactions in our crosses requires further work. Nonetheless, we note that if *H. p. butleri* and *H. p. sergestus* have differentially fixed derived alleles at opposing ends of the Z chromosome, one of these recombinants could represent the ancestral haplotype. For example, the high fitness of individuals bearing a Z*_BS_* chromosome could potentially be explained if it were ancestral, and thus compatible with many alleles at autosomal loci.

In contrast, Z*_SB_* individuals are notable for high variance in fertility (Fig. 5D). They show a negative correlation between fertility and the proportion of their autosomes that is homozygous for *butleri* alleles (Fig. 6A), and the variance in their fertility can be explained largely by an interaction between chromosome 8 and chromosome 20 (Table 1, Fig. 6B). Females that are either homozygous or heterozygous at both loci are fully fertile, but individuals homozygous at one locus and heterozygous at the other are less fertile (Fig. 6D). As such, it is unclear whether this pair of loci have any effect on the sterility of F1 females, even though they clearly have some effect in the backcross we studied here.

### 4.2 Candidate genes and comparison with *Drosophila* hybrid incompatibility loci

Oocyte development fails in sterile hybrid females in *H. pardalinus* at Lepidoptera stages 3-4 (Fig 3) (homologous with stages 8-9 of oogenesis in *D. melanogaster*), a period characterised by border follicle cell migration (Yamauchi & Yoshitake, 1984). Within the Bayesian credible intervals of the sex-linked QTL that interacts with QTLs on chromosomes 8 and 20, we identified 24 transcripts differentially expressed between sterile and fertile females with orthologs known to be involved in oogenesis in either *D. melanogaster* or the Speckled Wood butterfly (*P. aegeria*). Three of these are known to be associated with border follicle cells.

Within the proximal epistatic Z-linked QTL at 4.65 cM, we identified only one candidate gene, *magu*, mutants of which are known to cause defective border cell migration in *D. melanogaster* (Raza *et al*., 2019). Within the distal Z-linked QTL at 55.08 cM, 8 candidate genes were identified. One of these, *Epidermal growth factor receptor* (*Egfr*), guides dorsal migration of border cells during *Drosophila* oogenesis stage 9 (Duchek & Rørth, 2001), and is also expressed in the ovarian transcriptome of *P. aegeria* (Carter *et al*., 2013). We found 11 candidates involved in oogenesis within the QTL on chromosome 20. One of these, the multi-PDZ domain protein *bazooka* (*baz*), regulates border cell migration (Pinheiro & Montell, 2004), is expressed in the *P. aegeria* ovarian transcriptome, and furthermore is notable for being highly overexpressed in sterile individuals (log_2_ fold change =-5.52 for transcript HMEL016161g1.t3, the sixth lowest value in the dataset [Fig. 8A]). On chromosome 8, HMEL037834g1.t2, with ortholog *Neurexin 1* (*Nrx-1*), stood out as having the third highest log_2_ fold change (5.51) in the dataset. While not known to be involved in oogenesis, *Neurexin 1* is known to influence expression of *gurken* (*grk*) (Geng & Macdonald, 2007). The asymmetrical localization of *gurken* mRNA is key for its function during oogenesis, to establish anterior-posterior and dorso-ventral axes in the egg and embryo, and *gurken* encodes a TGF*α* family signaling ligand that activates the intracellular MAP kinase pathway via the the product of *Egfr*.

Differentially expressed transcripts located within QTL intervals, such as those discussed above, represent candidate regions for *cis*-acting differences between the two subspecies. Investigation of differential expression on its own, we can also identify putative trans-acting effects, or downstream consequences of the QTLs identified here. *Trailer-hitch* (*tral* has strong differential expression, high overall expression in ovaries, and is known to be involved in *Drosophila* oogenesis at stages 8-9 (Fig. 8, Fig. S9) (Wilhelm *et al*., 2005; Snee & Macdonald, 2009). Like *Nrx-1*, *tral* is involved in specifying the localization of the dorsoventral patterning gene *grk*.

We also noticed that alternative splices of transcript HMEL015815g1, orthologous to gene *spire* (*spir*) stood out as outliers in Fig. 8A. Although mapping to chromosome 1 and not in a QTL, HMEL037834g1.t2 was significantly underexpressed in sterile individuals (log_2_ fold change = 4.85), and HMEL015815g1.t6 significantly overexpressed (log_2_ fold change = -3.94). *spire* is involved specifically in stages 8-9 of oogenesis in *D. melanogaster*, where it affects the dorsal-ventral and anterior-posterior axes of the egg (Dahlgaard *et al*., 2007; Wellington *et al*., 1999).

The genus *Drosophila melanogaster* has long been used as a model to study developmental genetics, including the genetic basis of hybrid sterility. Some classical Dobzhansky-Muller incompatibilities have been identified and characterized in the genus (Brideau *et al*., 2006; Tang & Presgraves, 2009; Bayes & Malik, 2009). Because *Drosophila* has XY sex determination, in hybrids it is normally males that show sterility (Haldane, 1922). However, hybrid female dysgenesis has been observed in *D. melanogaster* in so-called P-M hybrids, in which oogenesis arrests at a very early stage (Kidwell *et al*., 1977; Schaefer *et al*., 1979; Bingham *et al*., 1982). This phenotype is due to a loss of control of P element transposition, normally suppressed via the *Drosophila* piRNA pathway in P strain flies (Evgen’ev *et al*., 1997; Kelleher *et al*., 2012). Superficially, the *Heliconius* sterility phenotype described in this study parallels this *Drosophila* case. The hypothesis that transposon silencing through the piRNA pathway is mis-regulated in sterile female hybrids has been explicitly tested in a different *Heliconius* hybrid system, *H. melpomene* and *H. cydno*. A subset of transposable elements were indeed derepressed in F1 hybrids, but there was no evidence that piRNAs themselves or three proteins involved in the piRNA pathway were misexpressed (Pinharanda, 2017). In our case, low fertility *H. pardalinus* female hybrids expressed three proteins in the piRNA pathway (*piwi/aubergine*, *AGO2/3*, and *vasa*) at lower levels than in more fertile individuals, though only *vasa* expression differences were significant (Fig. S9). In addition, one of our candidate genes, *tral*, forms a complex with piRNA proteins that inhibits P element transposition of a variety of transposons Liu *et al*. (2011). A *Drosophila*-like transposon derepression mechanism is therefore plausible, but the evidence remains inconclusive at present.

### 4.3 Evolution of hybrid incompatibilities

*Heliconius p. sergestus* is endemic to the dry forests of upper Huallaga valley in the Andes, and is separated from *H. p. butleri* in the Amazonian lowlands by the intervening Cordillera Escalera (Fig. 1). Nonetheless, the two subspecies are known to come into contact occasionally, and some putative wild hybrids exist (Michel Cast *pers. comm.*; Brown, 1976; Rosser *et al*., 2019). Theory predicts that in the face of gene flow, DMIs are more likely to be maintained when they are linked to traits involved in divergent adaptations (Bank *et al*., 2012), and *Heliconius* provide a possible example of this (Merrill *et al*., 2011). Divergent selection to different habitats could thus have facilitated the evolution of sterility within *H. pardalinus*, in a similar fashion to hybrid inviability evolving between plant populations as a pleiotropic consequence of adaptions to heavy metals (Macnair & Christie, 1983).

However, an alternative hypothesis is that hybrid sterility arose during an initial split between *H. elevatus* and *H. pardalinus*, only to be lost by hybridisation between sympatric populations in the Amazon, but retained in the allopatric subspecies *H. p. sergestus*. Concatenated whole genome phylogenies are consistent with this: *H. pardalinus* is paraphyletic, with *H. p. butleri* more closely related to the widespread Amazonian species *H. elevatus* than to *H. p. sergestus* (*Heliconius* Genome Consortium, 2012; Rosser *et al*., 2019). Moreover, despite strong assortative mating, *H. p. butleri* and *H. elevatus* are known to be fully fertile, while crosses between *H. p. sergestus* and *H. elevatus* are sterile, with phenotypes similar to those found here between *H. p. sergestus* and *H. p. butleri* (Rosser *et al*., 2019). Intriguingly, in the central 250 kb region of high *F*_ST_ between *H. p. sergestus* and *H. p. butleri* (Fig. 7C), we observed a reduction in *D*_xy_ between the Amazon taxon *H. p. butleri* and *H. elevatus* (Fig. 7B). The notable drop in diversity (*π*) in this same region in both *H. p. butleri* and *H. elevatus* (Fig. 7A) suggests a strong, recent selective sweep that also introgressed between these sympatric populations. Given that this region is in the middle of the main Z chromosome QTL for sterility between *H. p. butleri* and *H. p. sergestus*, introgression of this region is a candidate for explaining the lack of hybrid sterility between *H. elevatus* and *H. p. butleri* in the Amazon.

## 5 Conclusions

The genetics of Haldane’s Rule and Dobzhansky-Muller incompatibilities have been extensively studied in *Drosophila* and a few other male heterogametic systems, but hitherto there has been little genomic work on female heterogametic systems. Our work with *Heliconius* butterflies represents the first such study in Lepidoptera. We employ thousands of markers across the genome to map multiple regions involved in hybrid female sterility, and show an especially large effect of the Z chromosome. By intersecting these results with the list of differentially expressed genes among fertile and sterile hybrids, we identify six candidate genes (*magu*, *Egfr*, *baz*, *Nrx-1*, *tral*, and *spir*) potentially involved in hybrid sterility. Many questions remain unanswered, and functional genetic studies will be required to understand the mechanisms of ovariole development failure in hybrids. Nonetheless, we were able to show that several of the major findings from studies of Haldane’s Rule in *Drosophila* male sterility (e.g., multilocus effects, epistasis, involvement of the sex chromosome) are replicated in female sterile hybrids in a female heterogametic system. Future work can now address the genetic basis of sterility, as well as the potential tie-in with selfish genetic elements and with genes that act to defend the genome against their replication.

## 6 Data Availability Statement

The data that support the findings of this study will be made openly available on a public database following acceptance of the article.

## 7 Acknowledgements

This work was funded by NERC grant NE/K012886/1 to KKD and Harvard University. We also thank SERFOR, the Peruvian Ministry of Agriculture, and the Área de Conservación Regional Cordillera Escalera for collecting permits (0289-2014-MINAGRI-DGFFS/DGEFFS, 020-014/GRSM/PEHCBM/DMA/ACR-CE, 040–2015/GRSM/PEHCBM/DMA/ACR-CE). We are extremely grateful to the following people for help and support with field work in Peru: Ronald Mori Pezo, Corita Cordova, Mario Tuanama, Jarreth Caldwell, Christian Pérez, César López, Stephanie Galluser, and Gerardo Lamas. We are also very grateful to Pasi Rastas for guidance using LepMap-3, and to Michael Turelli and Daven Presgraves for helpful discussions regarding QTL results. We thank Shreeharsha Tarikere and Wendy Valencia-Montoya for commenting on the manuscript.

## Supplementary information

### Linkage Map

For the reads aligned to Hpar, the linkage map comprised 159,952 markers (a 29% improvement on Hmel2.5, with a total map length of 1157.27 cM). Marey maps plotting physical distances against genetic distance showed that linkage maps created using Hmel2.5 aligned reads and Hpar aligned reads were broadly similar (Supplementary Figs. S4 and S5). However, with Hpar some additional large regions with low recombination were apparent (for example, on the distal end of chromosome 15). Plots of estimated recombination rates between all pairs of markers for both linkage maps showed no evidence of misplaced markers (Supplementary Fig. S6). To validate our genotypic data and linkage map, we performed a QTL analysis on a wing colour pattern trait (presence / absence of yellow apical dots on the forewing). As expected, this showed a significant QTL peak encompassing the gene *cortex* (Supplementary Fig. S7), which is known to be involved in yellow colour pattern elements in *Heliconius* (Nadeau et al. (2016) The gene *cortex* controls mimicry and crypsis in butterflies and moths. Nature 534, 106–110).

**Table S1:** Numbers of markers on each chromosome and chromosome length in centimorgans for linkage maps using reads aligned to Hpar (left) and Hmel2.5 (right) reference genomes.

| Chromosome | Hpar |  | Hmel2.5 |  |
| --- | --- | --- | --- | --- |
|  | n markers | cM | n markers | cM |
| 1 | 11627 | 54.84 | 9353 | 53.57 |
| 2 | 4923 | 55.98 | 3537 | 55.98 |
| 3 | 5558 | 49.52 | 4342 | 46.57 |
| 4 | 4153 | 50.2 | 3038 | 49.04 |
| 5 | 5698 | 60.46 | 4618 | 57.29 |
| 6 | 8715 | 51.22 | 6853 | 50.03 |
| 7 | 8383 | 48.84 | 6914 | 48.84 |
| 8 | 5820 | 59.53 | 4549 | 53.66 |
| 9 | 5395 | 62.06 | 4193 | 50.32 |
| 10 | 11760 | 55.84 | 8840 | 55.87 |
| 11 | 6682 | 51.8 | 5333 | 50.44 |
| 12 | 9825 | 57.4 | 7818 | 55.93 |
| 13 | 10806 | 49.1 | 8278 | 48.95 |
| 14 | 4570 | 54.94 | 3452 | 54.96 |
| 15 | 6474 | 44.27 | 4919 | 45.46 |
| 16 | 6037 | 88.75 | 5041 | 69.48 |
| 17 | 10856 | 57.01 | 8272 | 57.04 |
| 18 | 9377 | 57.04 | 7427 | 53.81 |
| 19 | 10134 | 53.57 | 7847 | 53.63 |
| 20 | 8637 | 37.64 | 6616 | 38.68 |
| Z | 4527 | 57.29 | 3216 | 57.4 |
| TOTALS | 159957 | 1157.27 | 124456 | 1106.95 |

**Table S2:**
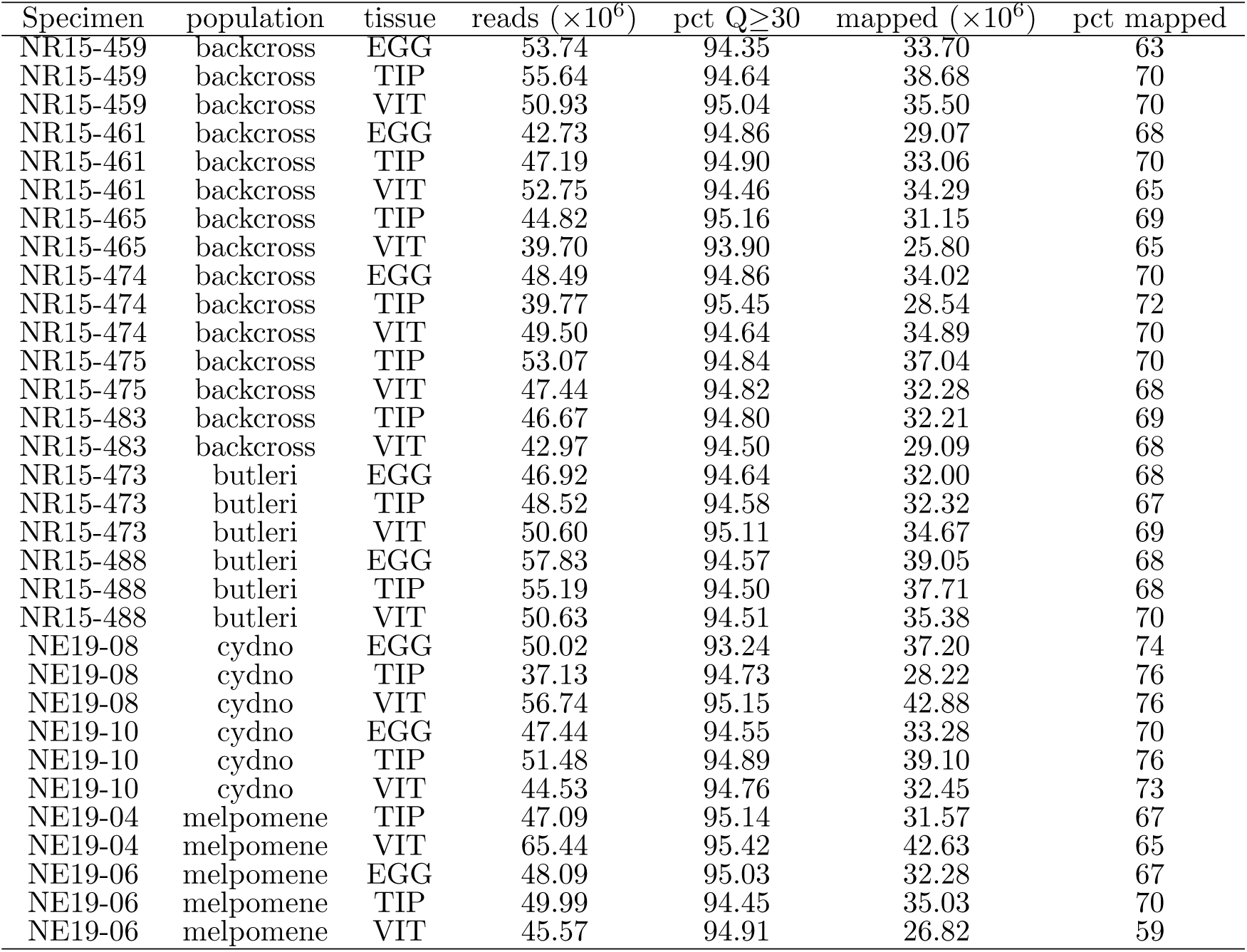
RNA sequencing and read mapping statistics to the Hmel2.5 reference.

**Table S3:**
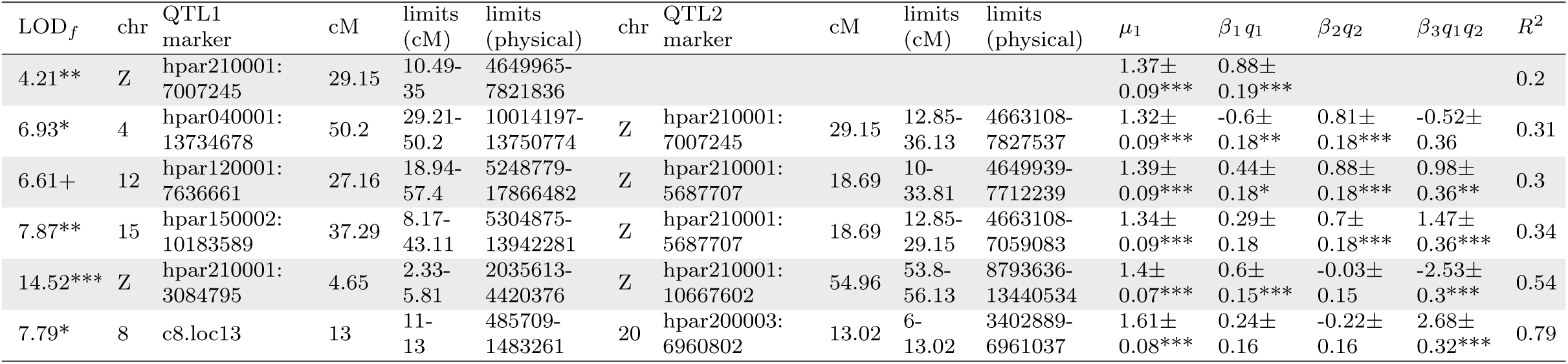
Summary of significant single locus (H_1_) and two locus (H_f_) QTL models using reads aligned to Hpar. The first column gives the LOD value of the full model (H*_f_*), with the significance estimated by permutation (+P*<*0.1, *P*<*0.05, **P*<*0.01, ***P*<*0.001). The next columns are the chromosome and QTL marker (scaffold and median physical position within the peak). The centimorgan limits are the Bayesian credible intervals, and the physical limits are the nearest typed flanking markers of that interval (all physical limits were on the same scaffold as the QTL peak, except for the chromosome 4 interaction † with Z, which was on scaffold Hmel204003 of Hmel2.5). The final five columns give the parameter estimates and *R*^2^ of the model. *β*_1_*q*_1_ and *β*_2_*q*_2_ are the estimated additive effects for the QTLs, i.e. the difference between the phenotypic averages for the alternative genotypes, and *β*_3_*q*_1_*q*_2_ is the coefficient for the interaction between the 2 loci. Model coefficients comprise the estimated value, the standard error, and the significance (thresholds as above). The significant interaction between chromosome 8 and 20 was detected using individuals holding a Z*_SB_* chromosome only.

| $LOD_f$ | chr | QTL1<br>marker | cM | limits<br>(cM) | limits<br>(physical) | chr | QTL2<br>marker | cM | limits<br>(cM) | limits<br>(physical) | $\mu_1$ | $\beta_1 q_1$ | $\beta_2 q_2$ | $\beta_3 q_1 q_2$ | $R^2$ |
| --- | --- | --- | --- | --- | --- | --- | --- | --- | --- | --- | --- | --- | --- | --- | --- |
| 4.21** | Z | hpar210001:<br>7007245 | 29.15 | 10.49-35 | 4649965-7821836 |  |  |  |  |  | 1.37±<br>0.09*** | 0.88±<br>0.19*** |  |  | 0.2 |
| 6.93* | 4 | hpar040001:<br>13734678 | 50.2 | 29.21-50.2 | 10014197-13750774 | Z | hpar210001:<br>7007245 | 29.15 | 12.85-36.13 | 4663108-7827537 | 1.32±<br>0.09*** | -0.6±<br>0.18** | 0.81±<br>0.18*** | -0.52±<br>0.36 | 0.31 |
| 6.61+ | 12 | hpar120001:<br>7636661 | 27.16 | 18.94-57.4 | 5248779-17866482 | Z | hpar210001:<br>5687707 | 18.69 | 10-33.81 | 4649939-7712239 | 1.39±<br>0.09*** | 0.44±<br>0.18* | 0.88±<br>0.18*** | 0.98±<br>0.36** | 0.3 |
| 7.87** | 15 | hpar150002:<br>10183589 | 37.29 | 8.17-43.11 | 5304875-13942281 | Z | hpar210001:<br>5687707 | 18.69 | 12.85-29.15 | 4663108-7059083 | 1.34±<br>0.09*** | 0.29±<br>0.18 | 0.7±<br>0.18*** | 1.47±<br>0.36*** | 0.34 |
| 14.52*** | Z | hpar210001:<br>3084795 | 4.65 | 2.33-5.81 | 2035613-4420376 | Z | hpar210001:<br>10667602 | 54.96 | 53.8-56.13 | 8793636-13440534 | 1.4±<br>0.07*** | 0.6±<br>0.15*** | -0.03±<br>0.15 | -2.53±<br>0.3*** | 0.54 |
| 7.79* | 8 | c8.loc13 | 13 | 11-13 | 485709-1483261 | 20 | hpar200003:<br>6960802 | 13.02 | 6-13.02 | 3402889-6961037 | 1.61±<br>0.08*** | 0.24±<br>0.16 | -0.22±<br>0.16 | 2.68±<br>0.32*** | 0.79 |

**Figure S1:**
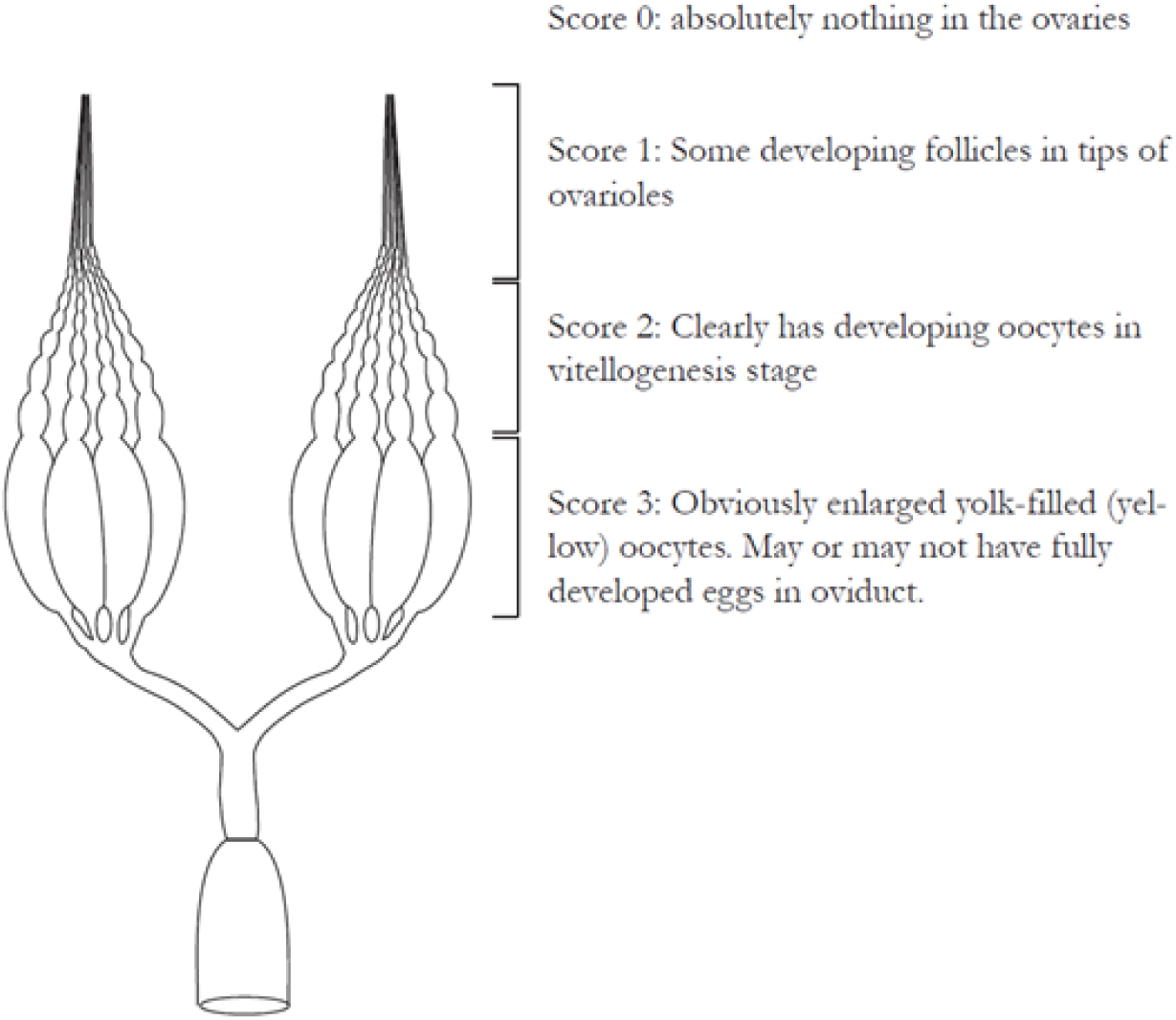
Fertility score guide. This scheme was used to characterize hybrid ovarioles in terms of gross developmental phenotype (Gullan & Cranston, 2014).

**Figure S2:**
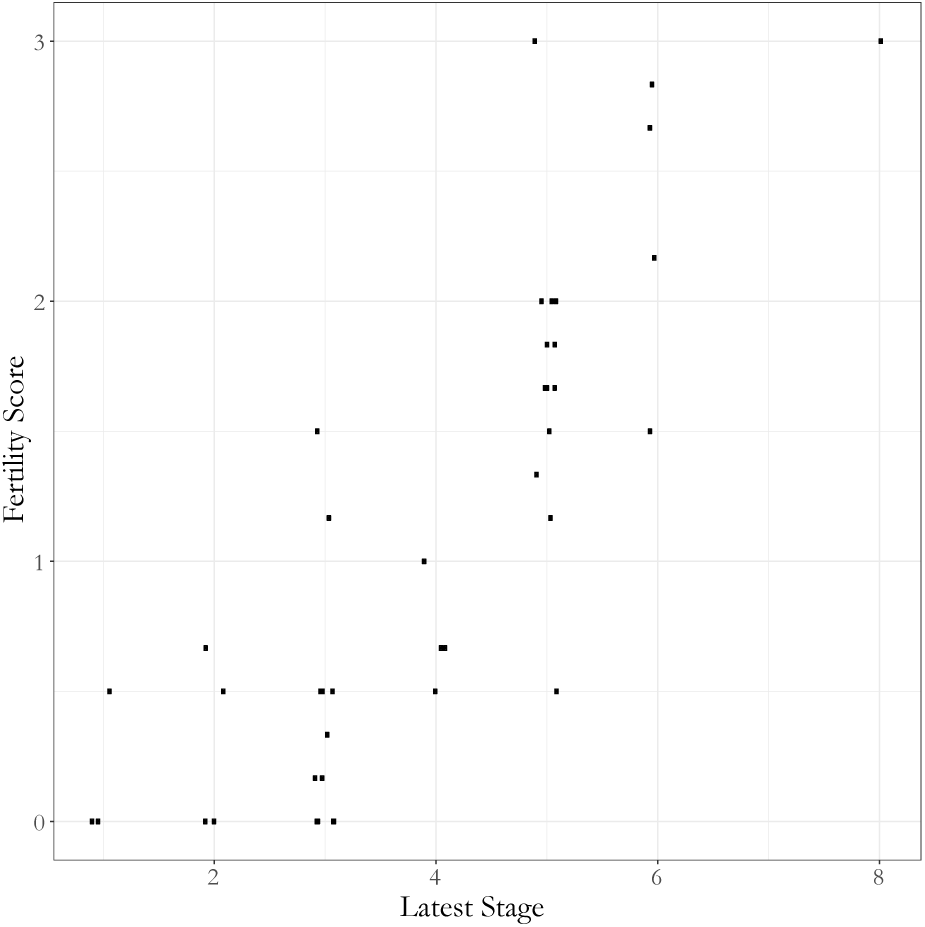
Comparison of scoring schemes. Among individuals for whom we were able to score both latest developmental stage and fertility score, the values were highly correlated. Note that the “Latest Stage” measurement is the latest stage observed in the sampled ovariole. Fully-developed eggs (stage 12, (Yamauchi & Yoshitake, 1984)) may have been present in the oviduct but were not observed in ovarioles themselves.

**Figure S3:**
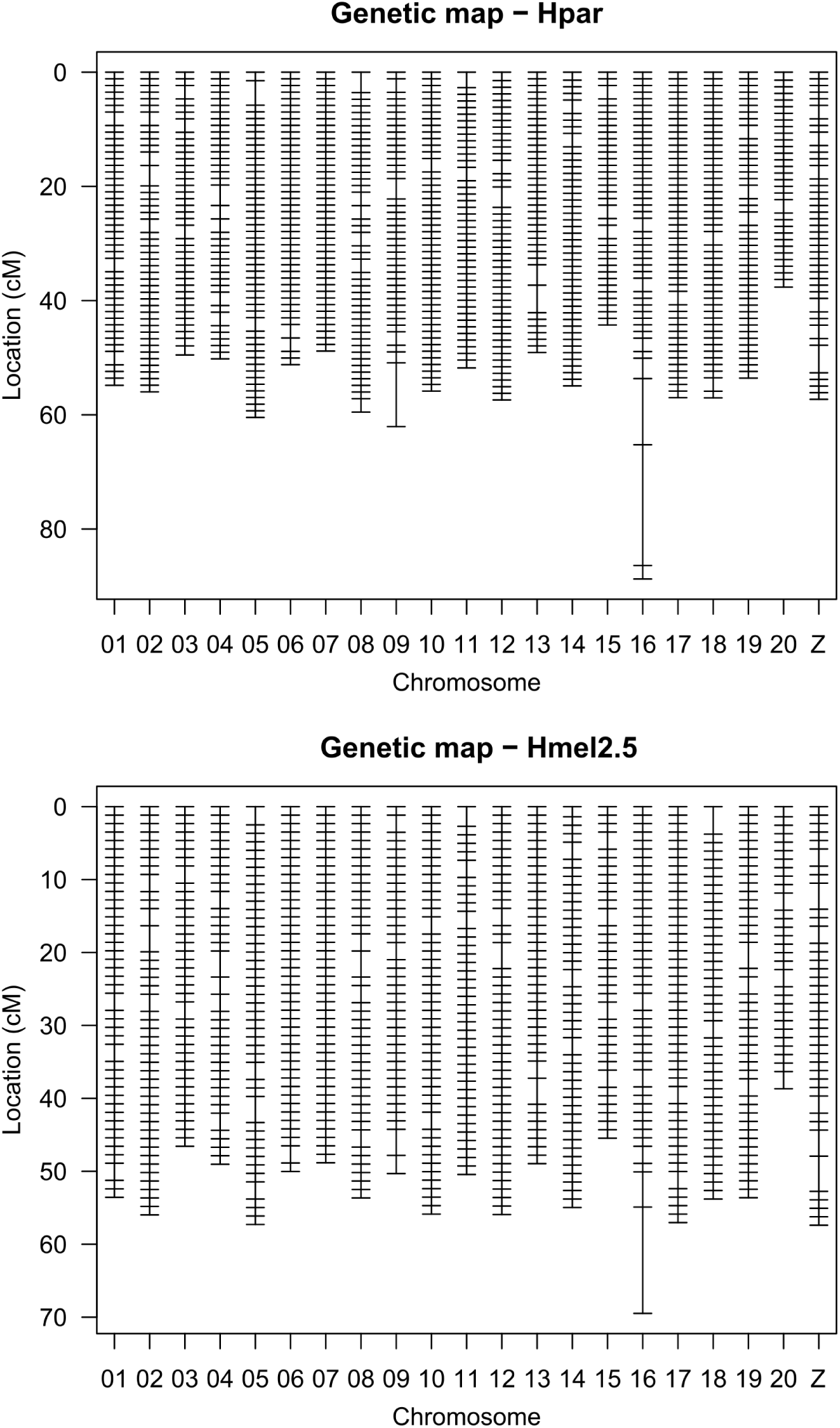
Marker locations for linkage maps using reads aligned to Hpar and Hmel2.5.

**Figure S4:**
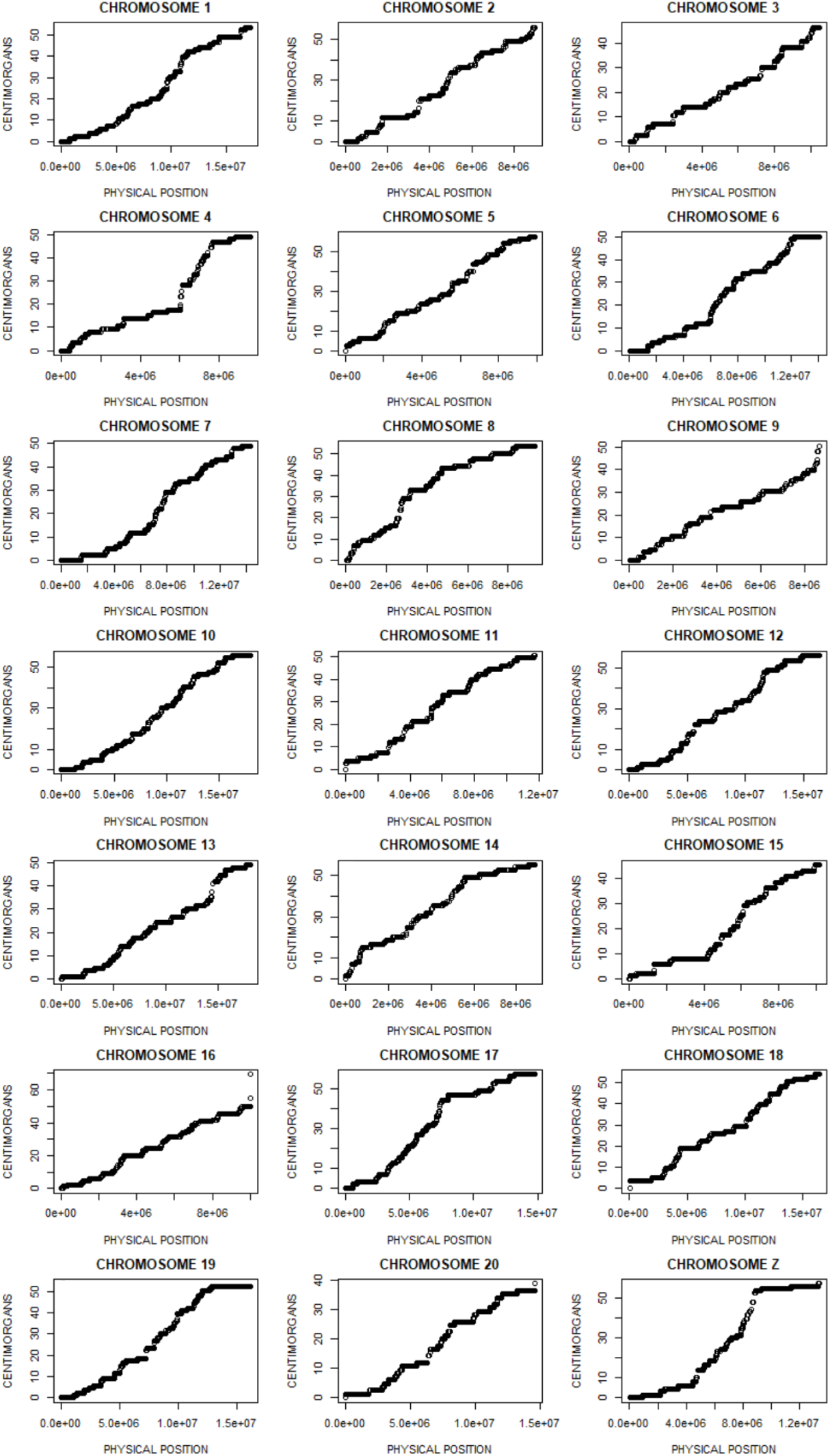
Marey maps using reads aligned to Hmel2.5.

**Figure S5:**
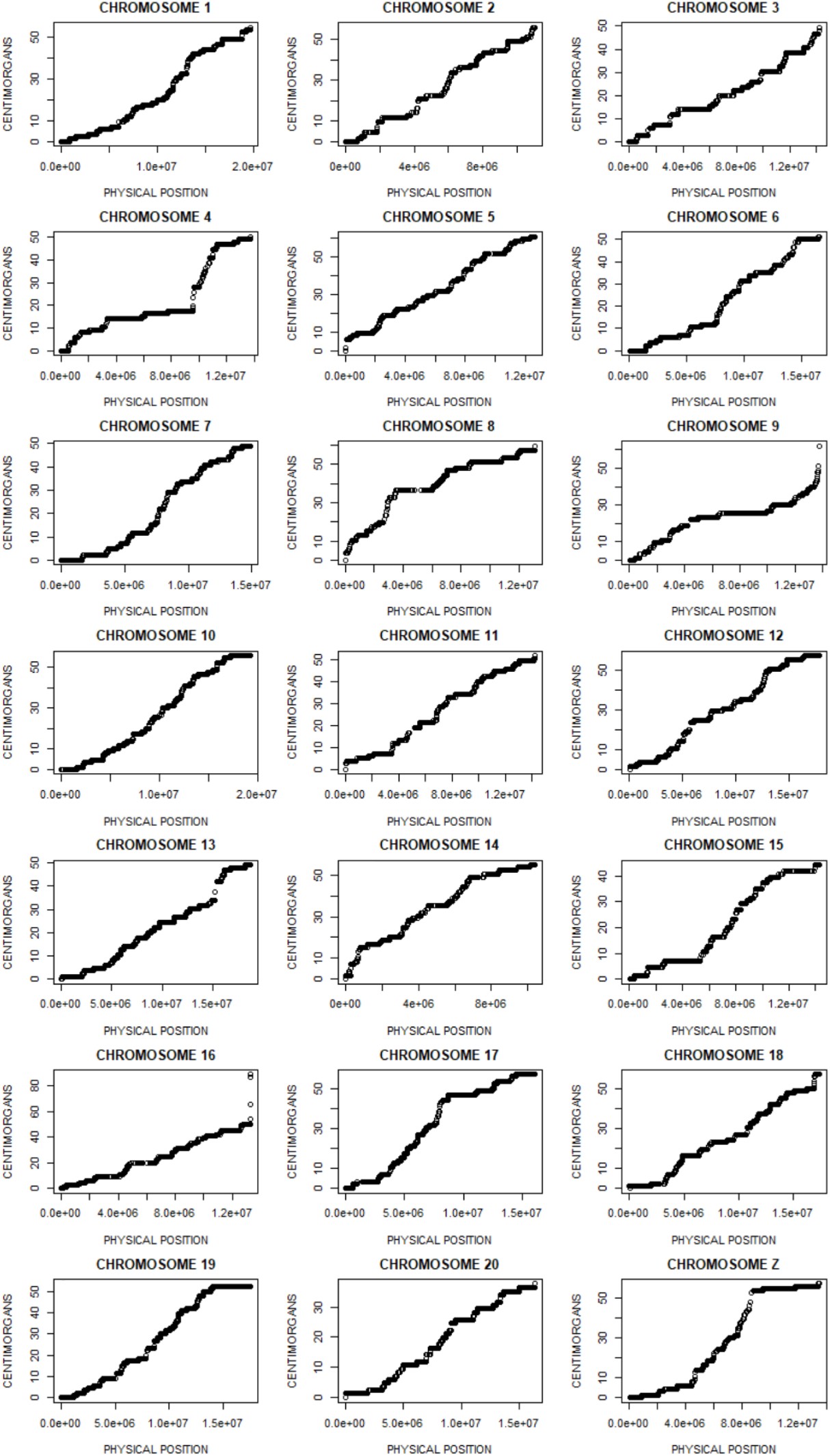
Marey maps using reads aligned to Hpar.

**Figure S6:**
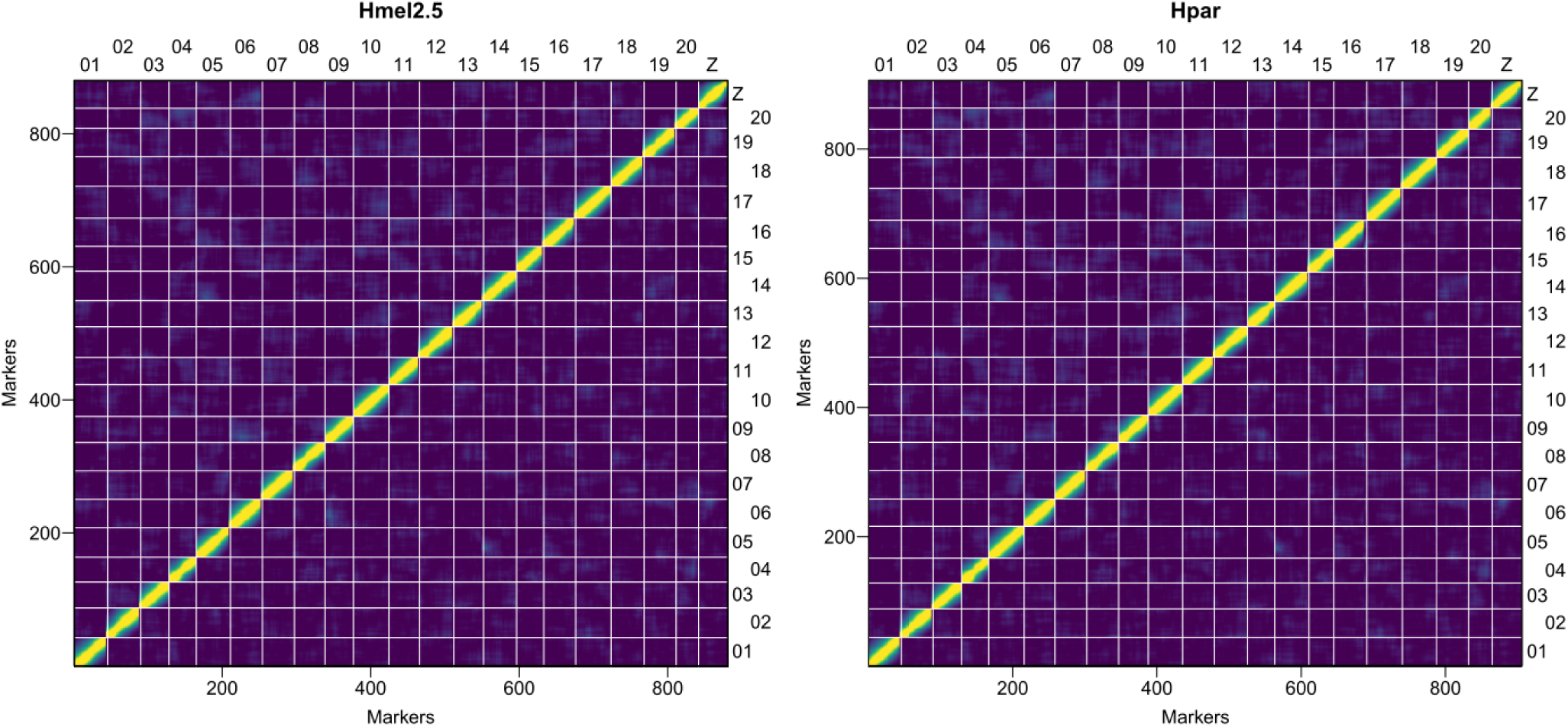
Pairwise recombination fractions (upper left) and LOD values for the test of recombination rate = 0.5 (lower right), using the Hmel2.5 and Hpar linkage maps. Yellow indicates linkage, blue indicates no linkage.

**Figure S7:**
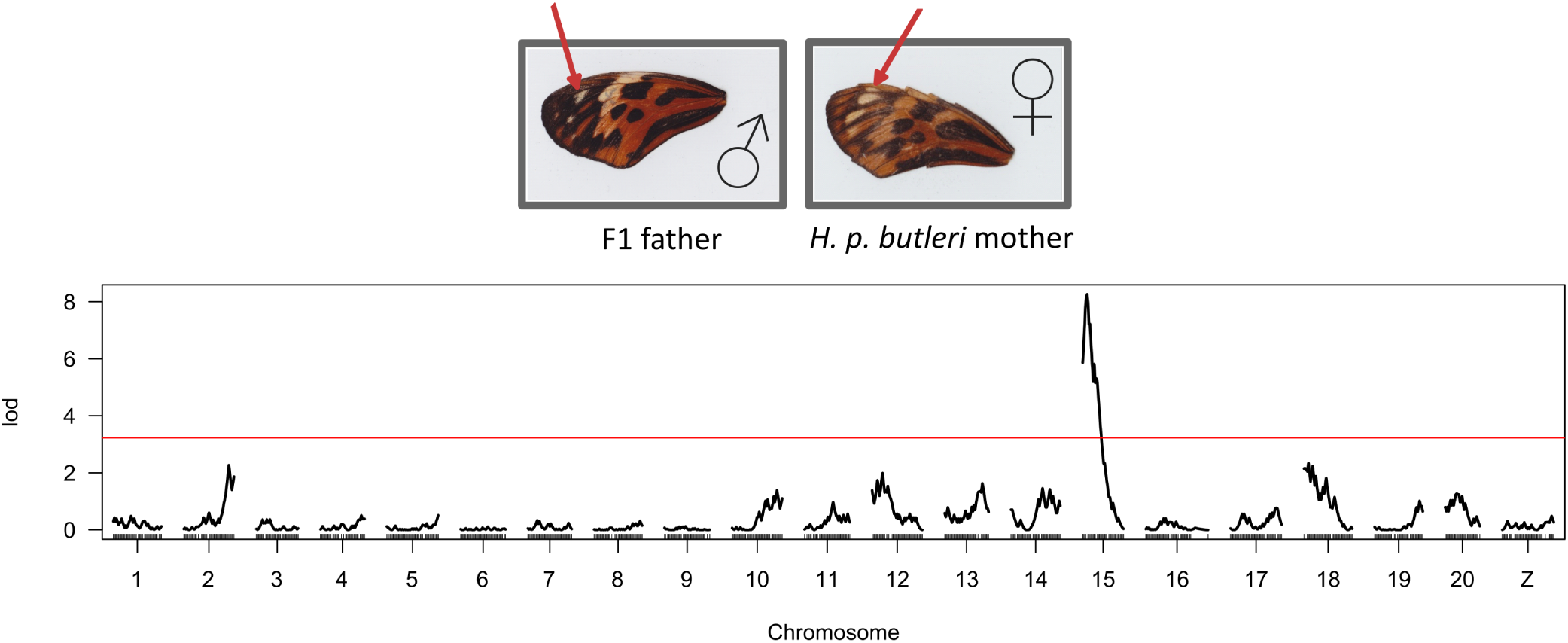
Colour pattern QTL. Backcross individuals produced by crossing a male F1 with a *H. p. butleri* female where scored by eye for large/reduced apical dots on the forewing (indicated by the red arrows in the figure). This trait was then analysed using Haley-Knott regression, and showed a significant QTL on chromosome 15, in the region of the gene *cortex*.

**Figure S8:**
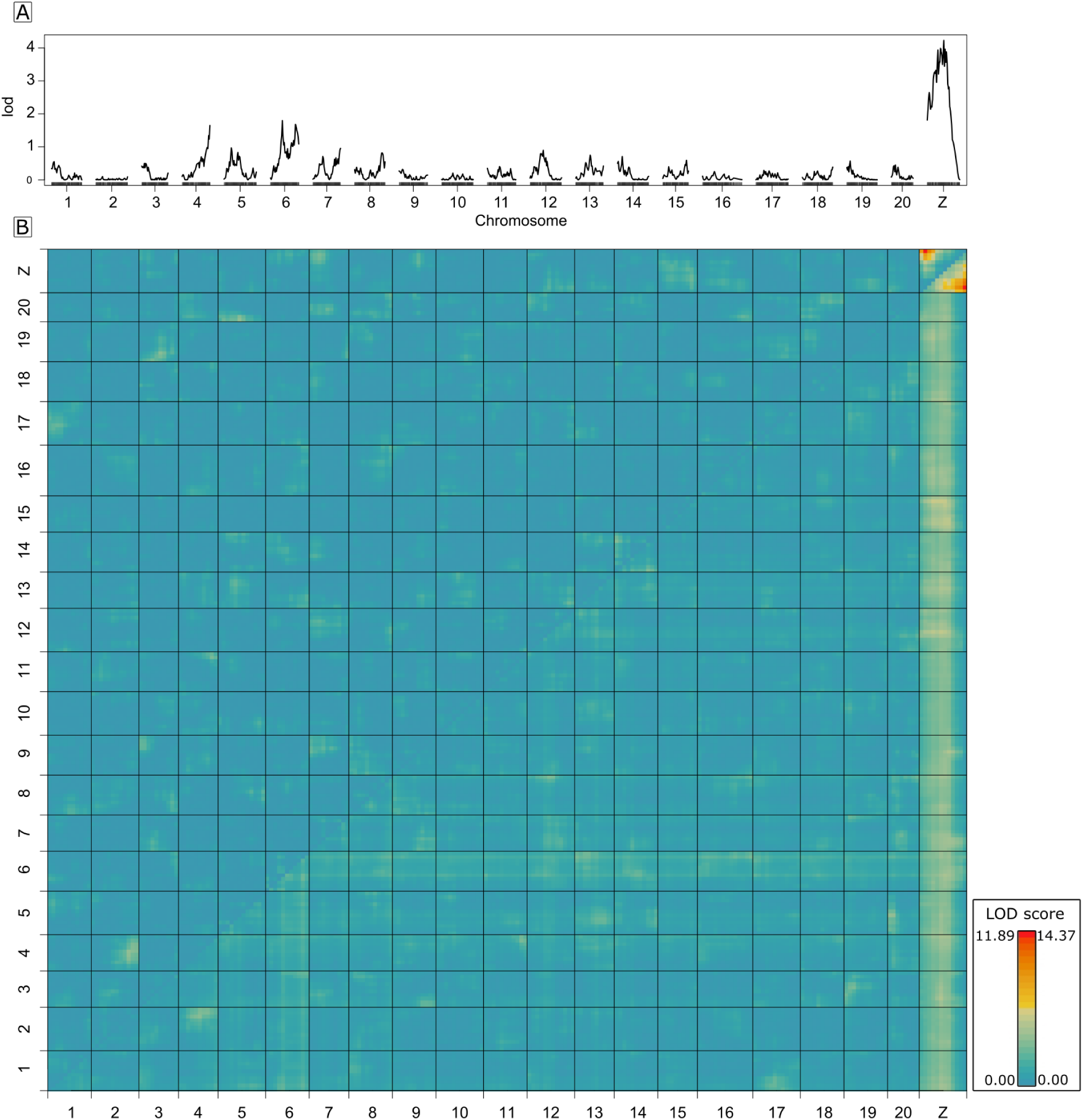
QTL genome scans controlling for kinship, using reads aligned to Hmel2.5. **A.** LOD values for fertility score as a function of genotype. **B.** Heat map for LOD*_f_* values (upper right triangle) and LOD*_int_* values (lower left triangle) between pairwise combinations of markers across the genome. Blues indicate low values, reds indicate high values. Max LOD *_int_* is left of the colour ramp, and max LOD*_f_* is to the right of the colour ramp.

**Figure S9:**
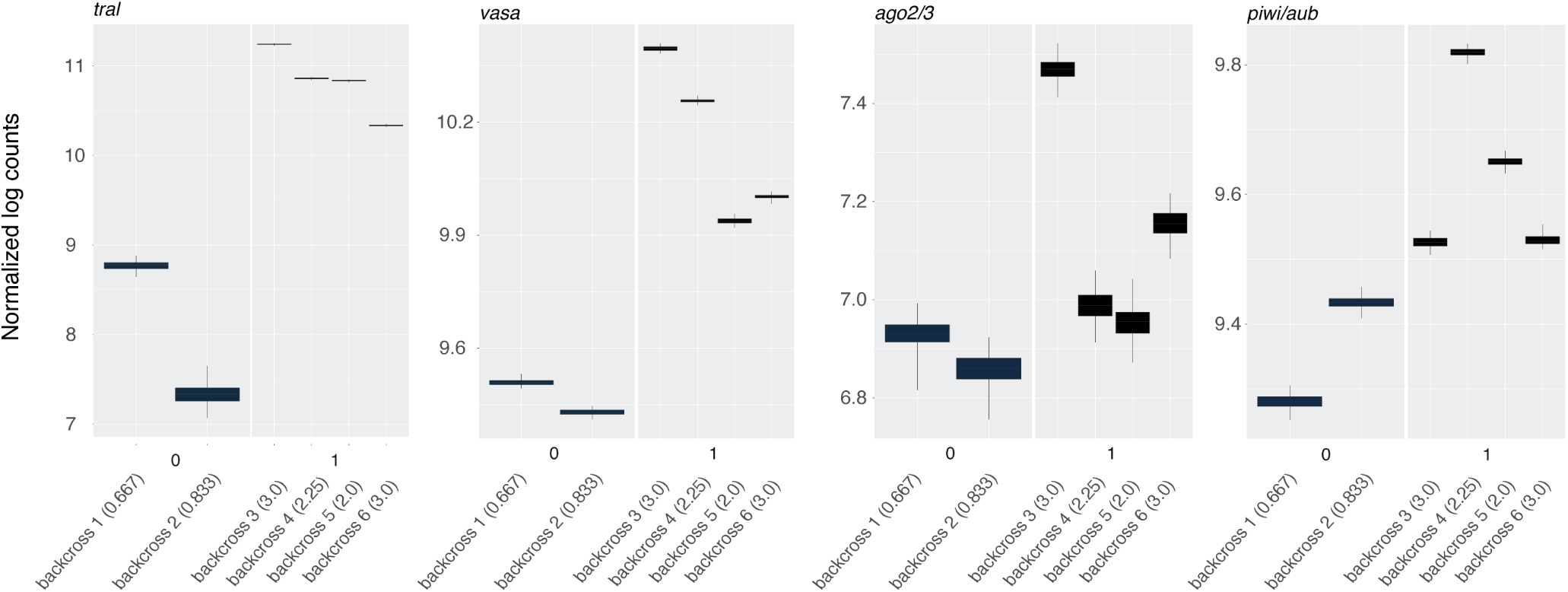
mRNA expression of piRNA-related transcripts. Expression levels for *tral*, *vasa*, *AGO2/3*, and *piwi/aubergine* are shown. Individuals are categorized based on whether they contained vitellogenic follicles (1) or not (0). In addition, each individual’s fertility score is indicated in parentheses. In each case, abundance is lower in undeveloped ovaries, but only significantly so for *tral* and *vasa*. Width of bars represents the interquartile range of bootstrap-resampled reads for each individual. Lower whisker is the smallest observation greater than or equal to lower edge of bar - 1.5 * IQR. Upper whisker is the largest observation less than or equal to upper edge of bar + 1.5 * IQR.

